# Network structure of the mouse brain connectome with voxel resolution

**DOI:** 10.1101/2020.03.06.973164

**Authors:** Ludovico Coletta, Marco Pagani, Jennifer D. Whitesell, Julie A. Harris, Boris Bernhardt, Alessandro Gozzi

**Affiliations:** Functional Neuroimaging Laboratory, Center for Neuroscience and Cognitive Systems @ UniTn, Istituto Italiano di Tecnologia, Rovereto, Italy; Center for Mind/Brain Sciences, University of Trento, 38068 Rovereto (TN), Italy; Allen Institute for Brain Science, Seattle, USA; Multimodal Imaging and Connectome Analysis Lab, McConnell Brain Imaging Centre, Montreal Neurological Institute, McGill University, Quebec, Canada

**Keywords:** fMRI, hubs, gradients, connectome, neuromodulation, Default Mode Network, co-activation patterns

## Abstract

Fine-grained descriptions of brain connectivity are fundamental for understanding how neural information is processed and relayed across spatial scales. Prior investigations of the mouse brain connectome have employed discrete anatomical parcellations, limiting spatial resolution and potentially concealing network attributes critical to the organization of the mammalian connectome. Here we provide a voxel-level description of the network and hierarchical structure of the directed mouse connectome, unconstrained by regional partitioning. We show that integrative *hub* regions can be directionally segregated into neural sinks and sources, defining a hierarchical axis. We describe a set of structural communities that spatially reconstitute previously described fMRI networks of the mouse brain, and document that neuromodulatory nuclei are strategically wired as critical orchestrators of inter-modular and network communicability. Notably, like in primates, the directed mouse connectome is organized along two superimposed cortical gradients reflecting unimodal-transmodal functional processing and a modality-specific sensorimotor axis. These structural features can be related to patterns of intralaminar connectivity and to the spatial topography of dynamic fMRI brain states, respectively. Together, our results reveal a high-resolution structural scaffold linking mesoscale connectome topography to its macroscale functional organization, and create opportunities for identifying targets of interventions to modulate brain function in a physiologically-accessible species.

## Introduction

Studies examining the structural architecture of the brain have advanced our knowledge of how information is processed and integrated across distributed and specialized neural circuits. Current network theory applied to brain connectomes has greatly contributed to this process, highlighting a series of common organizational principles underlying brain connectivity, many of which appear to be species and scale invariant ^1^. These include the presence of discrete regional sub-systems (termed *communities*) critically interlinked by a small number of highly-connected *hub* nodes, a configuration optimally suited for effective information processing and integration of neural signals across sensory and cognitive domains ^2,3^. Brain communities and hub regions have been observed at different investigational scales and using multiple connectivity readouts in several species, from the nematode *C. Elegans* to humans ^4–6^.

Recently, the mesoscale connectome of the mouse brain has been mapped via the use of directional viral tracers, representing one of the best characterized directed mammalian connectome ever described to date ^7–9^. The integration of this dataset with gene expression maps and layer-specific viral tracing have advanced our understanding of the wiring principles of the mammalian brain, revealing a network core of highly interconnected and metabolically costly hub nodes ^10^, and a phylogenetically conserved feedforward-feedback laminar hierarchy in intracortical structure ^8^. However, most investigations of the mouse connectome to date have been limited by the use of pre-defined anatomical parcellations in which connectional parameters, from which network attributes are computed, are quantified under the assumption of regional homogeneity ^7^. This has typically entailed the interrogation of subsets of anatomically aggregated meta-regions (for example, 213 x 213 regions in ^10^, or 130 x 130 in ^6^), an option that greatly increases the computational tractability of the mouse connectome. The use of predefined meta-areas is however non ideal, as the sharp inter-areal boundaries that characterize most neuro-anatomical parcellations reflect a discretization of otherwise regionally continuous cytoarchitectural or anatomical parameters which may straddle cross-regional network features. Moreover, the use of meta-regions greatly limits the spatial resolution of connectional mapping, potentially obscuring sub-regional attributes and motifs that could be critical to the network organization of the mammalian connectome. This in turns limits the potential of using the mouse as a species in which to probe salient nodal properties with sufficient regional precision.

Here we leverage a voxel-level data driven model of the mouse connectome ^9^, to provide a brain-wide, high-resolution (15,314 x 15,314 matrix, 270 um^3^) description of the network structure and hierarchical organization of the directed mouse connectome. Our results show that the mouse connectome is characterized by a finer network topography than previously reported, uncovering some previously underappreciated network features of the mammalian connectome. These include a segregation of hub regions into source and sink nodes, pointing at an organizational hierarchy in which higher order cortical areas serve as primary sources of neural output to the rest of the brain, and basal ganglia are configured as pivotal recipients of incoming projections. Using *in silico* network attacks, we also uncovered a strategic role of ascending modulatory nuclei as essential orchestrators of network communicability, a connectional property that makes these systems points of vulnerability for network function. We also found a tight inter-dependence between functional and structural brain organization, entailing the spatial arrangement of mouse cortical areas according to a hierarchy reflecting unimodal-transmodal and modality-specific functional processing, hence broadly reconstituting basic organizational principles of the primate brain. Our findings define a high-resolution structural scaffold linking mesoscale connectome topography to its macroscale functional organization, and create opportunities for identifying targets of interventions to modulate brain function in a physiologically-accessible species.

## Results

### Global hubs and rich-club core of the voxel-wise mouse connectome

A defining characteristic of brain connectomes is the presence of spatially localized set of integrative *hub* regions, characterized by high connectivity density ^2^. In order to identify regional features exhibiting hub-like properties at the voxel scale, we first mapped voxels exhibiting high connectivity strength using a spatially-resampled (15,314 x 15,314) version of the Allen Institute voxel-level mouse connectome ^9^, irrespective of the directionality of the connections. We termed the identified regions as *global hubs* to distinguish them from further hub identification carried out using the directed connectome (described below). This analysis revealed several sub-regional foci characterized by high connectivity strength (Figure 1A). The identified areas were prominently located in associative cortical regions such as the prefrontal, anterior cingulate, posterior parietal and retrosplenial cortices (Figure 1A). An additional large cluster of hub nodes was apparent in dorsal hippocampal areas. Our fine-grained mapping also allowed the recognition of a focal set of hub nodes encompassing focal portions of the basolateral and central amygdala. Notably, the spatial extension of the identified hub clusters in most cases encompassed only a marginal portion of the corresponding anatomical structure as defined in a high-resolution version of the Allen Brain Atlas (Figure S1A), suggesting that prior mapping of hub-like properties in the parcellated connectome might have been resolution-limited.

**Figure 1.**
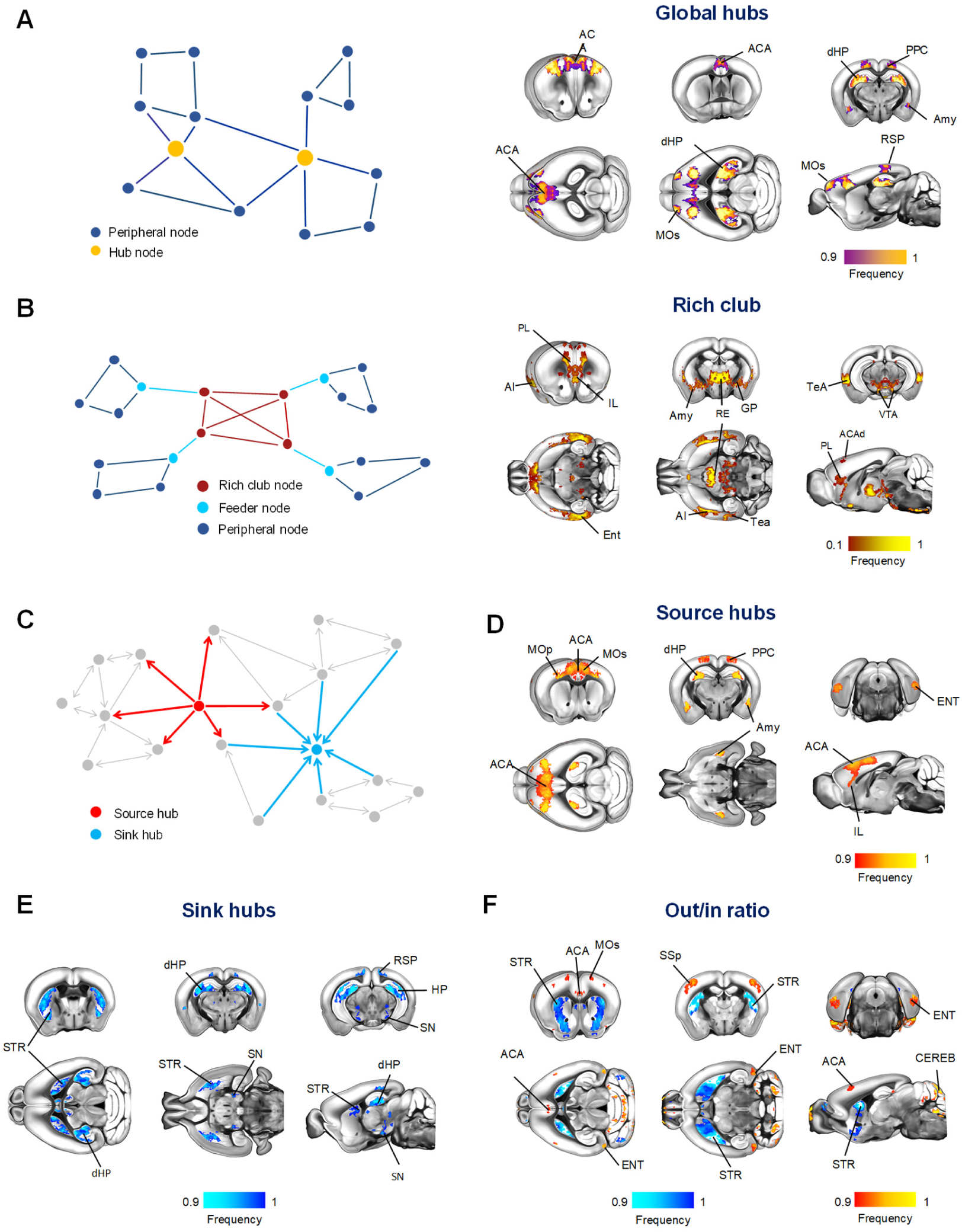
Source and sinks hubs of the mouse connectome are spatially segregable. A) Anatomical distribution of global hubs of the voxel-wise mouse connectome. Global hubs (yellow nodes on the left panel) were defined on the basis of nodal total strength. A frequency map was obtained by computing the fraction of times a node scored among the highest ranking strength nodes, limiting the visualization to the nodes that were classified as hubs at least 90% of the times. (B) Anatomical distribution of the rich club (red nodes on the left panel) of the voxel-wise mouse connectome. The frequency map indicates fraction of times high-degree nodes were retained as significant with respect to a set of random networks. (C) Network schematic illustrating our topological classification of high strength regions into neural sources (red) and sinks (light blue). Source (D) and sink (E) hubs were defined on the basis of the voxel-wise strength of outgoing and incoming connectivity, respectively. Frequency maps were computed as in (A). (F) Out/in ratio mapping. For each node, we computed the ratio between the strength of the outgoing and incoming connectivity. Frequency maps were obtained by computing the fraction of times a node scored among the highest (red/yellow) or lowest ranking (light blue/blue) nodes as in (A). Anatomical abbreviations are defined in Supplementary Table I.

Prior cross-species analyses of brain connectomes have revealed that highly interconnected regions define a core network structure, often referred to as *rich club*, which supports the efficient integration of otherwise segregated neural systems ^5,10,11^. To obtain a description of the mouse brain rich-club unconstrained by regional partitioning, we employed the procedure described by ^10^, benchmarking our mapping against 1,000 weighted rewired networks characterized by the same empirical in and out degree distribution ^12^. The obtained map revealed a more extended spatial topography than observed with global hub mapping, encompassing two major antero-posterior integrative axes (Figure 1B). The first of these included transmodal cortical integrators of sensory input (i.e. insula and temporal association cortex ^13^). The second axis encompassed infralimbic and mid-thalamic components of the fronto-hippocampal gateway ^14^. As observed with global hubs, most nodes of the rich club exhibited a sub-regional distribution with respect to a predefined high-resolution anatomical parcellation (Figure S1A). Notably, nodal mapping also revealed the participation of midbrain nuclei such as the ventral tegmental area, pointing at a previously unappreciated involvement of ascending dopaminergic nuclei as integral components of the rich club of the mouse connectome.

### Hub regions can be directionally segregated into neural *sinks* and *sources*

Our initial analyses were aimed at mapping global network features, and as such were carried out on a non-directed version of the mouse connectome. However, directional encoding can critically add key information to the topological organization ^15^, revealing organizational motifs undetectable in symmetrized connectomes. To probe how the direction of structural connections affect network attributes, we parsed high connectivity strength regions based on their directional profile, resulting in the identification of a set of segregable nodes which we termed *source* and *sink*, characterized by high-strength outgoing or incoming connections, respectively (Figure 1C).

Source node distribution broadly recapitulated the location of global hubs, encompassing higher order areas such as the anterior cingulate and posterior parietal cortices, amygdala, dorsal hippocampus, together with posterior entorhinal areas (Figure 1D). Interestingly, mapping of sink nodes revealed the involvement of dorsal hippocampal areas along with a new set of substrates, which comprised the basal ganglia throughout their antero-posterior extent (Figure 1E). Participation of nuclei within the *substantia nigra* was also apparent. These results show that high connection strength regions can be segregated based on their directional profile, and point at an organizational hierarchy in which higher order areas, such as the prefrontal cortex, serve as primary sources of neural output to the rest of the brain, while basal ganglia are pivotal recipients of incoming projections.

The observation of segregable sink and source high-connection strength areas prompted us to investigate whether such a hierarchy could be expanded to non-hub areas (i.e. to all brain regions, independent of their connection strength), by computing the voxel-wise ratio between outgoing and incoming connection strength, a metric which we term “out/in ratio” ^16^. This analysis would also allow us to differentiate regions characterized by a net connectional imbalance from those exhibiting both high input and output density (e.g. dorsal hippocampus).

The resulting out/in ratio map (Figure 1F) revealed a prominent configuration of basal ganglia as regions characterized by a low ratio of outgoing/incoming connections, corroborating a configuration of these substrates as connectivity *sinks*. Conversely, foci exhibiting a high out/in connection ratio were identifiable in higher order cortical areas, such as the anterior cingulate and entorhinal cortices, but also prominently encompassed some *non-hub* substrates, such as the cerebellum, and primary motor-sensory regions. Taken together, these results show that the directed connectome is topologically rich, and configured according to a global hierarchy that can be used to segregate regions in primary sources or receivers of axonal connections. In keeping with the above-described characteristics of hub and rich-club regions, the identified voxel clusters exhibited a clear sub-regional distribution with respect to a pre-defined high-resolution anatomical parcellation (Figure S1B).

### Structural communities of the voxel-wise connectome recapitulate large-scale fMRI networks of the mouse brain

The presence of segregable communities of tightly interlinked nodes is a hallmark feature of all mammalian connectomes mapped to date ^17^. Prior investigations of the community structure of the mouse connectome have entailed either metaregions ^6^ or have been limited to the sole cortical mantle ^8^. To identify stable brain-wide communities in the directed connectome with voxel-resolution, we used a multiscale modular decomposition approach ^6^ (Figure S2).

This approach revealed five prominent communities, encompassing different combinations of cortical and subcortical regions (Figures 2B and S3). Of note, the identified structural communities exhibited a spatial distribution closely recapitulating previously described resting state fMRI (rsfMRI) communities of the mouse brain ^18,19^. The first of such communities comprised trans-modal cortico-limbic areas as well as the dorsal striatum and antero-medial thalamus, spatially reconstituting key components of the mouse default-mode network (DMN ^20^). A second prominently cortical module encompassed motor-sensory latero-cortical, striatal and thalamic nuclei which have been previously described to compose a rsfMRI network termed *latero-cortical network* (LCN), which is functionally segregable and anticorrelated to the mouse DMN ^19^. A third module encompassed septo-hippocampal areas, while the fourth comprised olfactory areas and basal forebrain regions, once again recapitulating corresponding rsfMRI functional communities ^18^. Of note, anatomically similar structural connectivity partitions were also obtained using an agglomerative hierarchical clustering procedure (Figure S3) corroborating the validity of the nodal partitioning reported here.

**Figure 2.**
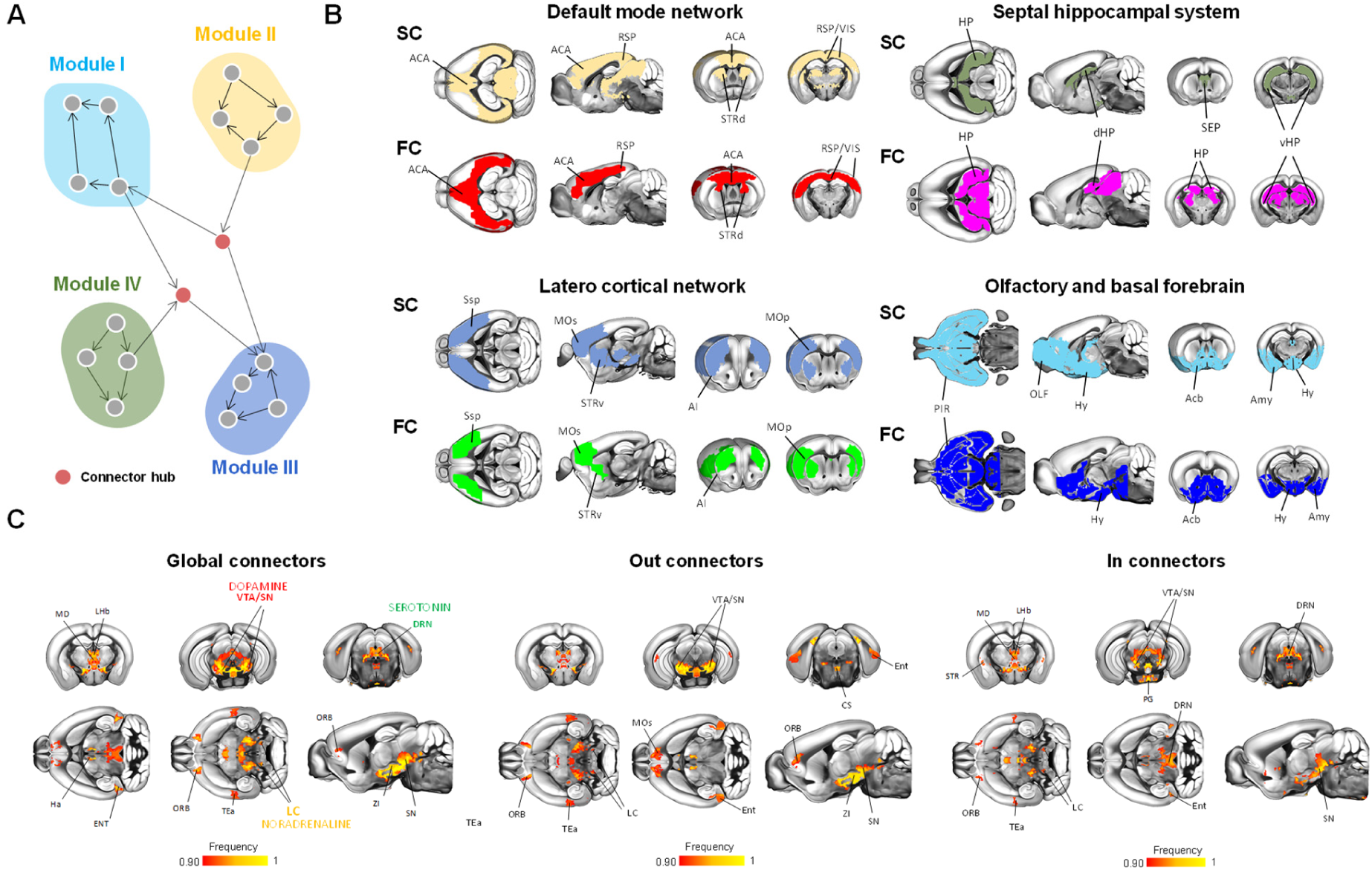
Connector hubs encompass key ascending neuromodulatory nuclei. (A) Network schematic illustrating a graph-based definition of communities and connector hubs of the directed mouse connectome. (B) Structural communities anatomically recapitulate functional (rsfMRI) networks of the mouse brain. Structural communities (top row) derived from a multiscale modular decomposition of the voxel-wise structural connectome (SC) were matched to corresponding functional communities obtained using a modular decomposition of rsfMRI functional connectivity (FC) (bottom row, from Liska et al., 2015). (C) Neuromodulatory nuclei are configured as connector hubs. Global (left), out (middle) and in (right) connector hubs were computed on the basis of the participation coefficient metric, accounting for outgoing or incoming connections only. Frequency maps were obtained by computing the fraction of times a node scored among the highest ranking nodes, limiting the visualization to the nodes that were classified as hubs at least 90% of the times. Anatomical abbreviations have been listed in Supplementary Table I.

Differences in the community structure of the axonal and functional connectome were also observed, an expected finding given the different nature and investigational scale of these two independent datasets, the lack of cerebellar coverage in the fMRI dataset examined, and the known resolution limit of modularity based algorithms ^21^. These discrepancies included the identification of a fifth cerebellar-pontine module in the structural connectome (SC), and the presence of a mid-thalamic community in the functional connectome (FC, Figure S4). Notwithstanding these discrepancies, the close resemblance between structural and functional communities supports a robust structural basis for the DMN, LCN and analogous distributed rsfMRI connectivity networks, a notion consistent with analogous investigations in human and primates ^22,23^.

Prompted by these correspondences, we probed the relationship between voxel-wise structural and functional connectivity (SC and FC, respectively) by carrying out a correlational analysis for the DMN, LCN and hippocampal network, three well-characterized distributed rsfMRI networks of the mouse brain ^24^. In keeping with recent investigations in primates ^25^, we found that voxel-wise correlation between FC and SC was non-linear, reflecting segregable connection length-dependent contributions (Figure S5). Specifically, functional-structural correlation was moderate to high (Spearman’s rho 0.35, 0.45, and 0.34 for DMN, LCN, and the hippocampal network, respectively) for relatively short connections (<1 mm, e.g. the scale of mouse cortical width), but lower for longer-range links (> 2mm, Spearman’s rho 0.26, 0.38, and 0.17 for DMN, LCN, and the hippocampal network, respectively). Interestingly, the correlation between FC and SC was instead robust and linear when both quantities were resampled at a lower spatial resolution using an anatomical parcellation (Pearson’s r, 0.59, 0.55, 0.54, p< 0.0001 for the DMN, LCN, and hippocampal network, respectively). This result is consistent with rsfMRI connectivity being a neural mass phenomenon reflecting the pooled activity of relatively large ensembles of neurons, exceeding the finer spatial scale of the voxel-wise mouse connectome ^26–28^.

### Ascending modulatory nuclei are configured as between-network connectors

The observation of segregable functional communities in the mouse connectome implicates the presence of *connector hubs*, i.e. nodal components critical to inter-modular communication. We first located connector hubs irrespective of the directionality of the connections, and termed the identified connector nodes *global connectors* (Figure 2C).

We found global connector hubs to be mainly localized in midbrain, hypothalamic and medio-dorsal thalamic regions, with only a marginal cortical involvement limited to orbitofrontal and temporal association areas. Remarkably, midbrain connector hubs focally encompassed three major set of ascending neuromodulatory nuclei, namely the ventral tegmental area and substantia nigra (dopamine), dorsal raphe nuclei (serotonin) and a set of voxels encircling the locus coeruleus (norepinephrine). Accounting for the directionality of the connections revealed evidence of a marginal topological segregation for some of the identified connector nodes (Figure 3C), the most noticeable difference being the configuration of periventricular (e.g. periaqueductal gray) and dorsal raphe nuclei as in-connectors. Of note, the high correspondence between functional and structural modules supports a role for connector nodes as strategic orchestrators of brain-wide network activity ^29^. In the light of this, our observations suggests that ascending modulatory system are strategically wired as key orchestrators of inter-modular activity, a finding consistent with emerging evidence pointing at a pivotal contribution of cathecolaminergic neurotransmission in modulating functional network activity and dynamics ^30,31^.

**Figure 3.**
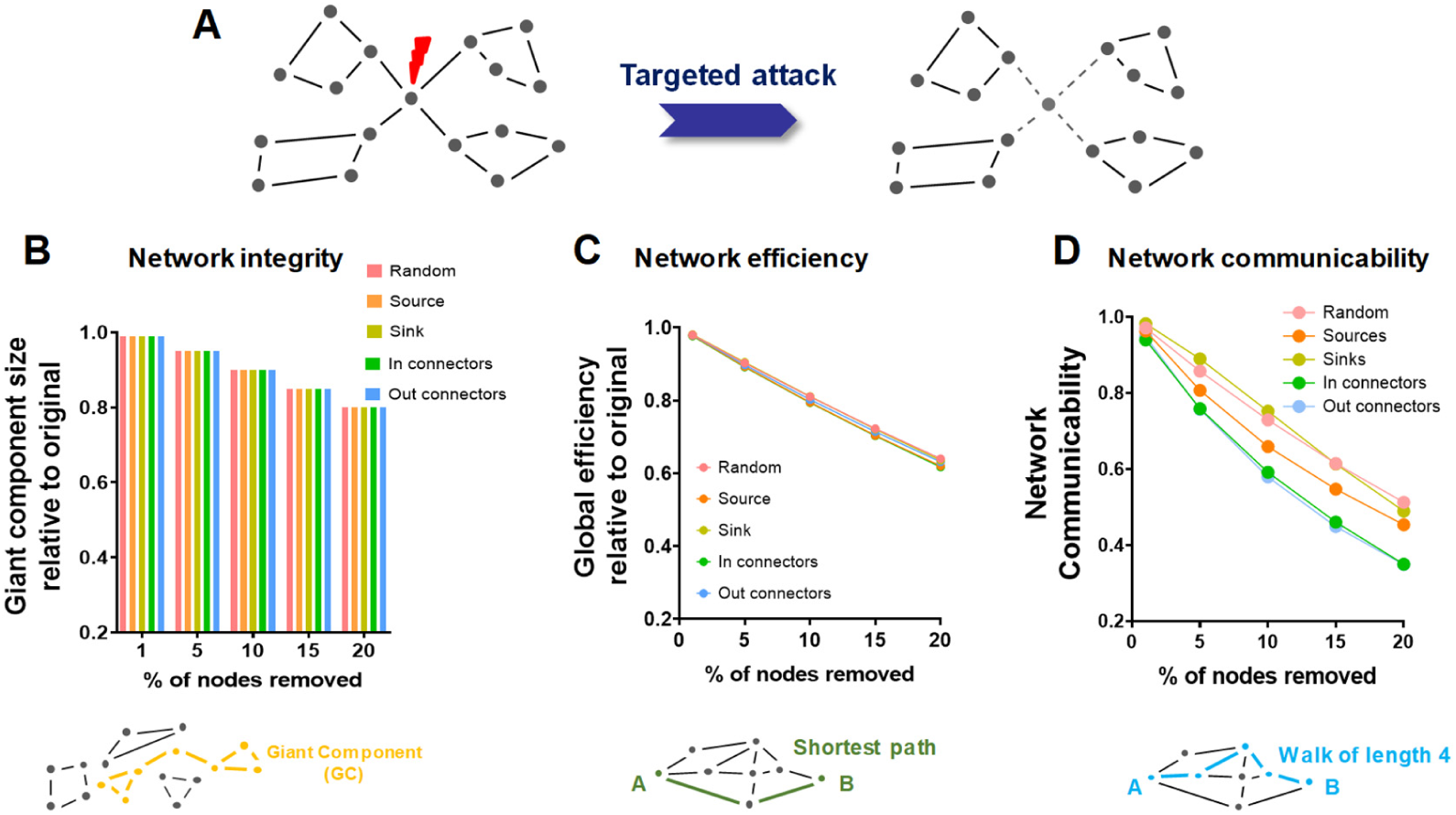
Connector hubs are critical effectors of network communicability. (A) Schematic illustration of targeted node removal, and its effect on network integrity. (B-D) Effect of targeted hub removal on different network properties. GC: giant component, GE: global efficiency; NE: network communicability.

### Connector hubs are critical mediators of network communicability

Graph theory postulates a critical contribution of hub regions to network integrity and stability, a notion supported by computational modelling of the human brain connectome ^32^. To test whether these assumptions hold for the voxel-wise mouse connectome, we performed a series of targeted in silico nodal attacks and assessed how these virtual lesions affect the ensuing network properties (Figure 3). The effect of hub (or random node) removal was assessed using two well-characterized global network attributes: (i) the size of the *giant component*, i.e. the largest subgraph in the network, a proxy for the network’s integrity ^32^ and (ii) global network efficiency, here measured as the average of the inverse of shortest path length ^33^.

Interestingly, targeted removal of sources and out-connector hub nodes did not produce appreciably larger network fragmentation than observed with random nodal attacks (Figure 3). Similarly, the removal of central nodes had an overall marginal impact in decreasing network efficiency, producing a fragmentation that was on average only ∼1.5% greater than random node removal (p<0.01, Figure 3). These findings suggest that, irrespective of their classification and directionality, hub nodes of the voxel-wise connectome are not critical for the integrity and efficiency of the network, a property that makes the mouse connectome highly resilient to targeted perturbations. We therefore next probed whether hub regions could be key to global network communication, a property that in structural networks can be assessed in terms of *total network communicability*, a metric reflecting the ability to effectively transfer information across the network ^34^. Notably, we found that deletion of connector hubs resulted in a robust reduction of network communicability when compared to random node removal (p<0.001, Figure 3). Taken together, these findings suggest that connector hubs, and the neurotransmitter nuclei therein contained, besides acting as pivotal modulators of between-module communication, are also configured as key effectors of global network communication.

### Gradients of structural and functional connectivity in the mouse cortex exhibit comparable topology

Recent investigations of the mammalian connectome have shown that the spatial arrangement of cortical connectivity in both human and primates reflects two superimposed gradients along which cortical locations are ordered according to their similarity in connections to the rest of the cortex ^23,35–37^. A first dominant cortical gradient is anchored in sensorimotor regions and radiates toward higher-order transmodal areas; a second separate gradient exhibits instead an axis of differentiation between sensorimotor modalities ^23,38^. Importantly, the organization of the unimodal-transmodal gradient is thought to define a network hierarchy of increasing functional integration which guides the propagation of sensory inputs along multiple cortical relays into transmodal regions, a view consistent with classic anatomical studies in primates ^39^.

To probe whether a similar organization is phylogenetically conserved in rodents, we applied diffusion map embedding ^40,41^ to the directed cortical connectome, with the aim to map gradients of connectivity profiles with voxel resolution (Figure 4). To account for the directional encoding of the connectome, the procedure was applied to a matrix mapping the connectional profile of each node, i.e. incorporating the information provided by both incoming and outgoing connections. We found that the structural connectome exhibits two spatial gradients of connectivity broadly recapitulating organizational principles observed in primates. Specifically, a dominant gradient (gradient A) involved a sensory-fugal transition between unimodal motor-sensory regions of the mouse LCN and transmodal components of the mouse DMN (Figure 4A). A second gradient (Gradient B) extended across unimodal visual and auditory cortices up to primary motor sensory regions, hence providing a regional differentiation between sensorimotor modalities.

**Figure 4.**
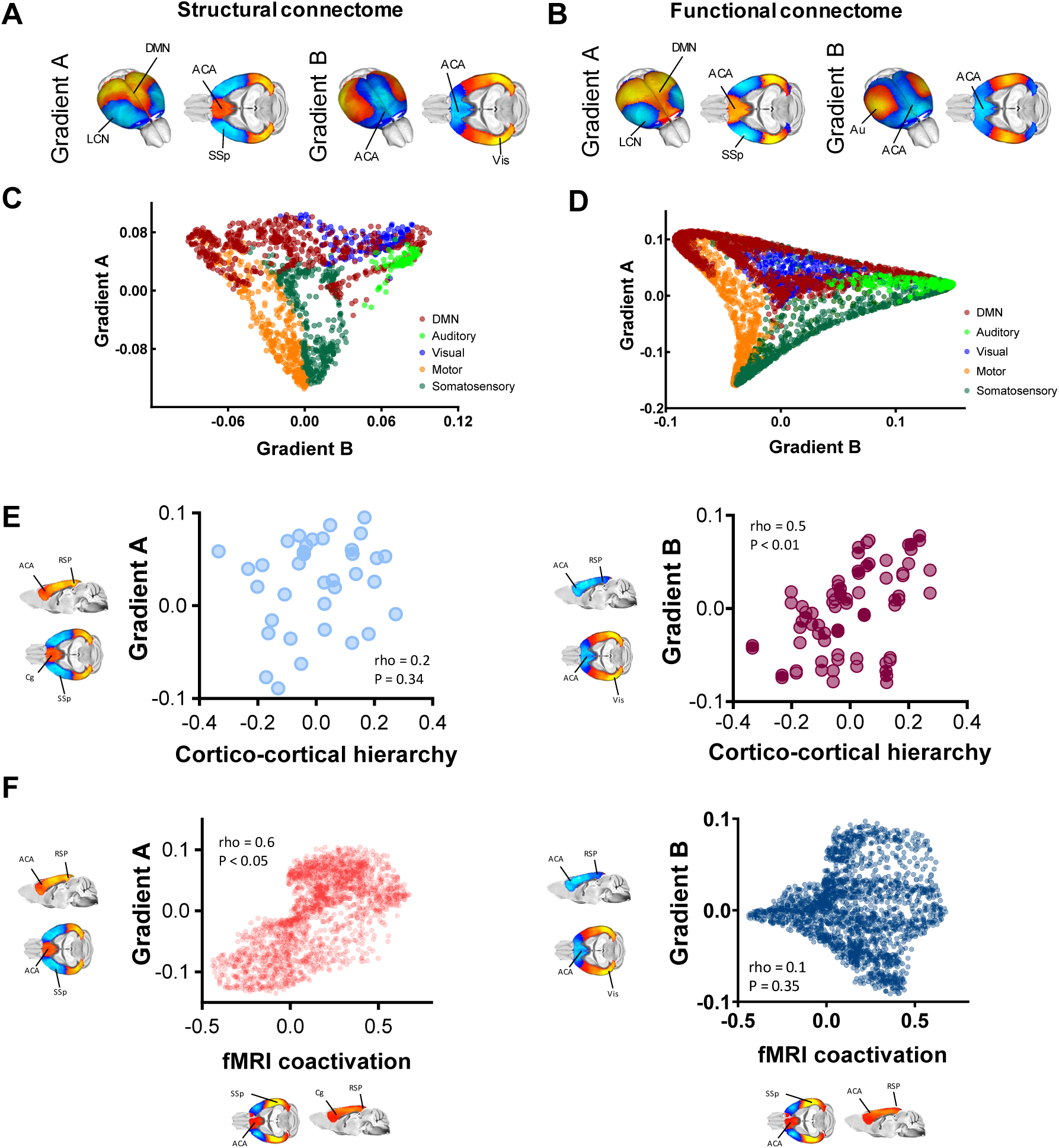
Gradients of structural and functional connectivity in the mouse cortex exhibit comparable topology. Structural (A) and functional (B) gradients of cortical organization in the mouse connectome. Gradient A encompasses a unimodal-polymodal spectrum of cortical regions extending from motor-sensory LCN (light blue/blue) to the DMN (yellow/red). Gradient B extends antero-posteriorly across primary sensorimotor (yellow) and transmodal associative regions (blue). (C-D) Regional scatter plots of gradient organization for SC (C) and FC. (E-F) Gradients of structural connectivity reflect cortico-cortical laminar hierarchy and constrain fMRI network dynamics. Modality-specific gradient B (Right), but not polymodal-unimodal gradient A (Left), reflects hierarchical intra-laminar organization of the mouse cortex. (B) Unimodal-polymodal DMN-LCN gradient A, but not modality specific gradient B, closely recapitulates the spatial topography of dominant cortical co-activation patterns governing fMRI dynamics in the mouse. Anatomical abbreviations have been listed in Supplementary Table I.

The close spatial correspondences between spatial organization of the functional and structural connectomes observed in our prior modular analyses prompted us to interrogate the presence of analogous gradients in the rsfMRI connectome (Figure 4B). Interestingly, brain-wide mapping of functional gradients revealed that the functional connectome is organized into two main cortical axes, including a unimodal-transmodal dominant gradient (DMN-LCN, Gradient A) and a modality-specific gradient (Gradient B, Figure 4B), hence spatially recapitulating key hierarchical features of the structural connectome. Spatial correlation of the two pairs of cortical gradients corroborated the presence of similar structural and functional topographies (Spearman’s rho = 0.83, p < 0.01 for gradient A and rho = 0.60, p < 0.05 for gradient B, p values corrected for spatial autocorrelation). An anatomical classification of the regional constituents of the identified gradients revealed that the topographical arrangement of trans-modal and unimodal areas was broadly comparable across modalities (Figure 4C-D), although a rearrangement in the spatial organization of modality-specific areas in functional gradient B was apparent, peaking in auditory-somatosensory regions as opposed to auditory-visual areas (Figure S6). Notwithstanding these modality-specific differences, the core architecture of cortical areas in the functional and axonal connectomes appeared to be largely similar, pointing at a common hierarchical organization for the functional and structural mouse connectome. Together, these results suggest that gradients of cortical organization in the mouse connectome recapitulate phylogenetically conserved architectural principles observed in higher mammalian species.

### Gradients of structural connectivity reflect cortico-cortical laminar hierarchy, and constrain fMRI network dynamics

Human studies have linked the organization of cortical gradients to hierarchical structure inferred from patterns of laminar cortical connectivity ^42^. The recent description of a feedforward-feedback laminar hierarchy in cortical regions of the mouse brain ^8^ allowed us to probe whether a similar organizational principle could explain the architectural organization of some of the gradients identified in the connectome. By computing the correlation between laminar hierarchy from Harris et al., (2019) and structural gradient topography in a set of corresponding cortical regions, we found that the regional organization of the modality-specific gradient (Gradient B) was robustly correlated with intracortical laminar hierarchy (Figure 4E, Spearman’s rho = 0.49, p<0.01, corrected for spatial autocorrelation). Intralaminar cortical hierarchy was instead not predictive of unimodal-polymodal gradient (Gradient A) topography (Figure 4E, Spearman’s rho = 0.17, p=0.33, corrected for spatial autocorrelation).

We however noted that the anatomical organization of the unimodal-polymodal gradient was anatomically consistent with dominant recurring patterns of fMRI co-activation (CAPs) recently described to govern spontaneous fMRI network dynamics in this species ^26,43^. These take the form of infraslow oscillatory transitions in latero-cortical and midline poly-modal regions reminiscent of the topographical organization of gradient A. In keeping with this notion, we found a strong spatial correspondence (Figure 4F, Spearman’s rho=0.60 and p<0.05, corrected for spatial autocorrelation) between gradient A and a mean functional CAP reflecting the dominant opposing relationship between the unimodal and transmodal areas governing spontaneous fMRI state dynamics. Conversely, the modal-specific gradient B did not show a significant relationship with the spatiotemporal structure of this dominant pattern of spontaneous brain activity (Figure 4F, right panel, Spearman’s rho = 0.09 and p = 0.35, corrected for spatial autocorrelation). These findings link mouse cortical gradient organization to patterns of laminar connectivity, and suggest that cortical hierarchy may critically shape the spatiotemporal structure of dominant patterns of spontaneous brain activity.

## Discussion

Here we provide a fine-grained description of salient architectural motifs of the mouse connectome, without the imposed limits of discrete regional parcellations. Departing from regional-constrained studies, we find that hub regions and core network components of the voxel-wise mouse connectome exhibit a rich topography encompassing key cortical and subcortical relay regions. We also typify regional substrates based on their directional topology into sink or source regions, and report a previously unappreciated role of modulatory nuclei as critical effectors of inter-modular and network communicability. Finally, we demonstrate a close spatial correspondence between the mesoscale topography of the mouse connectome and its functional macroscale organization, and show that, like in primates and humans, the mouse cortical connectome is organized along two major topographical axes that can be linked to hierarchical patterns of laminar connectivity, and shape the topography of fMRI dynamic states, respectively.

Our regionally-unconstrained mapping of hub-like regions complements and expands prior investigation of the mouse connectome, providing a spatially precise identification of network features and hierarchical motifs that may guide future manipulations of nodal properties in this species ^6^. These include a fine-grained localization of hub-like properties in sub-regional components of large integrative areas, such as the dorso-lateral hippocampus or the central and basolateral amygdala, which were previously been considered as regionally homogeneous ^6^. Similarly, our rich club mapping revealed a more detailed spatial topography than previously reported ^10^, revealing two major organizational axes of high relevance for sensory-integration and higher cognitive functions ^44–46^, and which recapitulate organizational features observed also in non-mammalian species ^47^. These results suggest that subcortical relay stations are core components of nodal rich clubs across evolution, serving as critical integrators between top-down and bottom-up functional processing.

Importantly, our results also revealed previously unappreciated organizational features of the mouse connectome that advance our understanding of the fundamental wiring principles of the mammalian brain in three main directions. First, the use of a high-resolution and directed connectome enabled us to segregate hub regions into *source* and *sink* areas. The ensuing classification revealed the emergence of a global hierarchy in which higher order cortical areas and hippocampal regions serve as primary sources of neural input to the rest of the brain, and basal ganglia (plus focal mesencephalic nuclei) are wired as major receivers of distributed neural input. This hierarchical configuration follows a phylogenetic gradient in the arrangement of structural connectivity, and is optimally designed for the execution of rapid motor responses in response to salient external stimuli ^48^. Such a hierarchical configuration could also be expanded to non-hub regions via a brain-wide computation of the ratio of outgoing and incoming connectivity strength, defining a related organizational axis with motor-related nuclei, such as the cerebellum and basal ganglia, being located at its extremes.

A second notable feature is our observation of a strategic configuration of ascending modulatory systems as connector hubs and essential effectors of network communicability. Previous investigations of the regionally-segregated mouse connectome have produced a largely cortico-centric description of connector hubs, involving cingulate, orbitofrontal and posterior association cortices, together with the basal ganglia and regionally undifferentiated midbrain regions ^6^. Our results shift the focus from the cortex to subcortical relay stations, and document that ascending neurotransmitter systems are central to the mouse connectome and are configured as inter-modular connector hubs. Importantly, the observed spatial correspondences between the structural and functional topography of the mouse connectome argue for a critical role for these neuromodulatory nuclei in shaping large-scale neural activity. This notion is consistent with the observation that catecholaminergic and serotonergic activity critically control functional network topography and dynamics ^30,31,49^. Together with the observation that connector hub removal critically diminishes network communicability, these results suggest that that ascending modulatory systems are strategically wired as central orchestrators of large-scale inter-modular communication, a network property that might be key in ensuring the effective and finely-tuned control of complex behavioral and physiological states exerted by these systems. At the same time these properties might render these nodes key points of vulnerability for functional network disruption in brain disorder, a notion consistent with emerging evidence linking neuromodulatory dysfunction to neurodegenerative pathologies ^50^. Interestingly, targeted removal of global hubs did only negligibly affect measurements of network integrity and efficiency when compared to random node deletion. This finding suggests that the mammalian connectome is structurally highly resilient, and argues against a role for this class of hub regions as critical mediators of network integrity in the mouse connectome. This result is partly supported by analogous investigations of the human connectome. For example ^51^ reported a linear decrease in global efficiency in a targeted attack for human structural networks, analogously to what we observed for the mouse connectome. Similarly, ^52^ found functional network fragmentation to occur only when about 75% of the high strength nodes were removed. It should however be noted that other reports seem to be at odds with these results, suggesting a significant vulnerability of the human connectome against targeted attacks in human networks (see ^32^ for a recent overview on the topic). Whether these discrepancies reflect modality- and resolution-related discrepancies, or a lower proportion of long-range integrative fibers in rodents owing to evolutionary scaling of white/grey matter ratio ^53^, remains to be established.

Finally, our voxel-wise description of two principal axes of cortical organization in the mouse connectome, and their topological linking with cortico-laminar organization and patterns of spontaneous fMRI dynamics, establishes a direct link between the mesoscale topography of the mouse connectome and its functional macroscale organization. These results suggest that the spatial arrangement of cortical areas along unimodal-polymodal and modality-specific gradients represents a general evolutionarily conserved principle governing the hierarchical organization of the mammalian cortex across evolution, and one that might intimately constrain the emergence and topology of spontaneous brain activity. This notion is consistent with a recent description of a cortical hierarchy in the parcellated mouse brain, as assessed by using an imaging marker of intracortical myelin content ^54^. Our findings expand these prior observations by providing cross-modal and voxel-wise evidence of two superimposing functional and structural cortical gradients broadly recapitulating organizational principles observed in the human and primate brain. These include a hierarchical organization reflecting a well-characterized feedforward-feedback laminar hierarchy ^8^, and a spectrum between unimodal regions and transmodal areas. It should be noted however that in the mouse the latter are known to exhibit a much lower degree of regional specialization than in primates, an observation that explains a categorization of latero-posterior visual and auditory territories as polymodal components of the posterior parietal cortex ^55,56^. Importantly, our results also revealed that a dominant cortical gradient spatially shapes the emergence of prevailing patterns of cortical co-activation governing spontaneous fMRI dynamics, further relating the topography of the connectome with the structure and temporal evolution of spontaneous cortical activity ^26^. The notion of a tight constraining effect of the structural connectome on functional network topography was further corroborated by evidence of largely overlapping functional and structural communities. This finding expands prior investigations of the mouse functional connectome ^57,58^, by highlighting a robust structural basis for distributed fMRI networks of the mouse brain such as the DMN and LCN. Such a close spatial overlap, however, does not appear to comprise hub topography, as previous voxel-wise mapping of functional hubs in the mouse only partly recapitulated the rich connectional features reported here ^18^. Such incongruity might reflect the fact that the spontaneous fMRI signal is a neural mass phenomenon, reflecting local and remote contributions that are negligibly constrained by more fine-grained topological features of the structural connectome.

In summary, here we provide a precise characterization of the network structure of the mouse connectome, with voxel resolution. Our results reveal a high-resolution structural scaffold linking mesoscale connectome topography to its macroscale functional organization, and create opportunities for identifying targets of interventions to modulate brain function and its network structure in a physiologically-accessible species.

## Material and methods

### Construction of the structural connectome

We estimated SC using a resampled version of the recently published voxel scale model of the mouse structural connectome ^9^ to make the original matrix computationally tractable. A whole brain connectome was built under the assumption of brain symmetry ^6,59^ resulting in a weighted and directed 15,314 x 15,314 matrix composed of 270^3^ µm voxels. Regional quantifications of network properties and correlations between structural and functional attributes were carried out using three main sets of predefined anatomical parcellations of the mouse connectome. To quantify the subregional localization of network attributes (Figure S1), we employed one of the finest parcellation available of the mouse connectome (i.e. the lowest hierarchical level in the Allen Mouse Brain Atlas, excluding layer-encoding ^8^). This parcellation was volumetrically matched to the sampling dimension of our voxels by discarding small nuclei whose spatial extension was – for either hemisphere – lower than the resolution of our voxel-wise connectome (45 regions out of 323, Table S2). Regional quantifications of sub-regional localizations were then limited to the remaining set of 278 areas. Correlation between functional and structural connectivity was carried out a set of meta-regions such to reduce spatial resolution and therefore maximize the contrast with corresponding correlations using voxel-wise sampling. The list of the 89 regions used for such comparisons is reported in Table S3. Meta-regions were selected such to cover the anatomical distribution of the functional modules described in ^18^. Regional quantification of structural gradient features and cortical hierarchy were carried out using the original cortical parcellation described in ^8^, corresponding to the isocortical subset in Table S2.

### Hub and rich club mapping

We computed normalized out-strength (source), in-strength (sink), and in+out strength (global) hub regions of the voxel-wise connectome. Hub regions were mapped by capturing the variability in the distribution of the metric of interest, via an iterative computation of the fraction of times a node scored among the top 50^th^ to 99^th^ percentile of highest ranking nodes, similarly to the “top percentage” approach outlined in ^60^ and employed in ^18^. We limited the visualization to the nodes that were classified as hubs at least 90% of the times, with the aim to capture top strength nodes. This approach ultimately led to the final representation of nodes exceeding the 94% strength percentile, for all hub categories of this manuscript.

The network core or “rich club” of the mouse connectome was mapped using with the weighted variant described in ^10^, limiting the analysis to the weighted ipsilateral connectome. Specifically, we computed the normalized rich club coefficient, defined as the ratio between the empirical rich club coefficient and the rich club coefficient obtained from an ensemble of 1,000 rewired networks where each network maintained the empirical in and out degree, together with the total wiring length of each node (as assessed by Euclidean distance ^12,61^). Due to the high computational demands of the rewiring procedure, we left a margin of 5% error on nodal wiring length constraint. Instead of testing all possible degree configurations, which usually range from 1 to *k* with k being the highest degree found in the network, we restricted the mapping between 6’720 and 8’143, corresponding to the 90^th^ and 99^th^ percentile, respectively, of the total degree distribution. This choice was motivated to both reduce the influence of low degree nodes, unlikely to represent hubs of the network, and to reduce the computational demands associated with rich club mapping with our high resolution matrix. Statistical significance (P < 0.05) was assessed by obtaining a P value directly from this null distribution. Across all normalized rich club coefficients, we computed for each node the fraction of times it was included in the rich club, similarly to the procedure described for the definition of source, sinks, and global hubs. We observed that all rich club coefficients tested in the above mentioned range yielded statistically significant results (p < 0.001).

### Multiscale modular decomposition and participation coefficient

We analyzed the network structure of the weighted and directed mouse structural connectome using the Louvain algorithm as implemented in the Brain Connectivity Toolbox ^33^. Similarly to the procedure outlined in^6^, we systematically varied (from 0.3 to 3.0 in 0.1 step, 100 repetitions at each step) the resolution parameters controlling the size of the modules, performing consensus clustering ^62^ and thus obtaining a representative community subdivision for each of the tested resolution setting. We then computed similarity between all the consensus partitions using the adjusted mutual information score, a similarity index ranging from zero to one, where zero means perfect disagreement and one means identical clusters. To avoid arbitrary thresholding of the similarity matrix, we performed again consensus clustering, limiting the threshold range from 0.9 to 1.0 (in 0.005 step), and obtaining at each step a representative hierarchical modular subdivision of the mouse structural connectome. We finally computed the agreement matrix, which encodes how many times any two hierarchical subdivisions of the connectome were grouped together. Figure S2 illustrates the procedure and shows the detection of stable modules across hierarchies. For all subsequent analyses, we focused on the first stable hierarchy level. Of note, using the normalized mutual information index (as in^6^) instead of the adjusted mutual information yielded similar results, with highest hierarchy level being identical across the two measures. We selected one resolution parameter among the ones constituting the highest hierarchy level, and at the selected spatial scale (gamma=1.0), we run 500 independent iterations of the Louvain algorithm, followed again by consensus clustering. We finally probed the statistical significance of the final partition against 1,000 randomly rewired networks characterized by the same empirical in and out degree distribution, and by maintaining the total wiring length of each node ^10,61^. Specifically, we used the total connectivity strength of each module as significant variable, reasoning that the internal cohesion of a given partition should be higher than expected by chance. We found that the total connectivity strength of each module always exceeded the total connectivity strength of the 1,000 rewired networks, suggesting that the degree sequences as well as the total wiring length of each node cannot adequately account for the spatial organization of the communities of the mouse structural connectome.

Module topography in the structural connectome was further corroborated using an agglomerative hierarchical clustering procedure of a matrix obtained by computing between-nodes similarity (as by Spearman Rank correlation) based on the connectivity profile of each node (see Figure S2). A comparison of the obtained clusters using the dice coefficient revealed an overall high concordance between the results obtained with these two procedures. We found a dice coefficient of 0.7 for the DMN, 0.82 for the LCN, 0.91 for the hippocampal module, and 0.92 for the olfactory-basal forebrain community. Finally, we found that the pontine-cerebellar module was almost equally represented by two clusters, one encompassing the cerebellum, and the other covering pons and medulla (dice coefficient of 0.66 and 0.56, respectively).

Our module detection procedure led to the identification of N = 7 modules, including two symmetric monohemispheric DMN and two olfactory-basal forebrain components, which we have joined into a single module for consistency with functional mapping and before computing their significance. The functional (rsfMRI) modules described in Figure 3 were obtained from ^18^. The procedure for functional module detection has been extensively described in the original work ^18^. To better match SC and FC modules, the basal forebrain and ventral midbrain modules identified in ^18^ were merged together to constitute a single ventral brain community. Finally, using the same procedure described for the definition of source, sinks, and global hubs, we computed the global (in+out), in- and out-connector hubs, ranking nodes based on their weighted participation coefficient, a measure of connection diversity ^33^.

### Virtual lesion mapping

The role of hubs for the network global functioning was probed by means of targeted virtual attacks. For each of the metrics of interest (in- and out strength, and global participation coefficient), we removed a given fraction of the highest ranking nodes (from 5 to 40%, in 5% step by zeroing all the incoming and outgoing connections), comparing the size of the giant component, global efficiency (measured as the average inverse shortest path length), and total network communicability, here limited to map the first indirect neighbors^34^. Metrics were computed pre- and post-attack, and changes with respect to these indices were expressed as a percentage of the intact network’s value. For each fraction of removed nodes, we compared targeted hubs deletion to 1000 random attacks, assessing statistical significance (P < 0.05) by obtaining a P value directly from the null distribution.

### Functional and structural gradients

We applied diffusion map embedding on SC and functional connectivity (FC) as previously described ^23,40,63^. Briefly, this nonlinear dimensionality reduction technique seeks to project high dimensional connectivity data into a lower dimensional Euclidean space, identifying spatial gradients in connectivity patterns. The cortical SC (FC) matrix is first mapped into an affinity matrix that represent the similarity of connectivity profiles across nodes. The eigenvectors describing the diffusion operator formed on the normalized graph Laplacian of the affinity matrix identify gradients in connectivity patterns over space.

To compute SC gradients, the structural affinity matrix was built based on the cortical connectional profile of each node, i.e. by incorporating the information provided by both incoming and outgoing connections. The functional affinity matrix was built using the same steps described by ^23^. In reporting the results we explicitly looked for gradients capturing the polymodal-unimodal differentiation as well as the modality specific organization of cortical connectivity as described in recent human and primate work ^23,64^. In line with human results, we found that the dominant gradient of both SC and FC recapitulates the unimodal-polymodal continuum (Gradient A in Figure 5A and 5B), whereas the second ranked functional and the third ranked structural gradients delineate comparable modality specific spatial configuration of cortical connectivity (Gradient B).

We additionally computed the correlation between SC gradients spatial maps and a dominant rsfMRI co-activation patterns (CAPS) published by ^26^, in an attempt to establish a link between the organization of the structural connectome and FC dynamics. In their work, ^26^ described three pairs of recurring oscillatory states account for the more than 60% of rsfMRI variance. Notably, two of these oscillating patterns are characterized by a conserved cortical topography entailing the opposing engagement of latero-cortical and DMN regions reminiscent of the mapped cortical gradients, the main difference between them being a differential involvement of subcortical structures (i.e. hippocampus). To correlate the topography of these dominant co-activation patterns with that of the structural gradients, we therefore generated a mean cortical CAP out of these two fluctuating states, using the mean value across the hemispheres. We did not consider the third pair of states (CAPs 3 and 4 in^26^) owing to its more widespread cortical topography and strong coherence with fMRI global signal, implicating the involvement of a possible global external input to the emergence of this meta-state. Finally, we also computed the correlation between SC gradients and cortical hierarchy scores computed on the basis of feedforward-feedback laminar connectivity patterns of the mouse brain as described and computed in^8^, using the same set of cortical brain regions described by the authors. For all the spatial correlational analyses involving gradients, we accounted for the spatial autocorrelation using Moran spectral randomization as implemented in the BrainSpace toolbox, using Euclidean distance between nodes as input for computing the Moran eigenvector maps ^40^.

### rsfMRI data

The rsfMRI dataset used in this work consists of N = 15 scans in adult male C57Bl6/J mice which are publicly available ^24,26^. Animal preparation, image data acquisition, and image data preprocessing for rsfMRI data have been recently described in greater detail elsewhere ^18,45^.

### Code and data availability

All graph theoretical analyses were performed using the Brain Connectivity Toolbox ^33^. All the other analyses were carried out using custom Matlab/Python scripts, FSL and ANTS, unless otherwise stated. Functional and structural maps, and the employed code will be made publicly available upon acceptance for publication of this manuscript.

## Acknowledgments

This work was funded by the European Research Council (ERC, DISCONN to A.G., Grant Agreement 802371). A.G. also acknowledges funding by the Simons Foundation (SFARI 400101, A. Gozzi), the Brain and Behavior Foundation (2017 NARSAD, Independent Investigator Grant 25861), the NIH (1R21MH116473-01A1), Telethon Foundation (GGP19177) and the University of Padua inter-departmental project Proactive. M.P. is supported by European Union’s Horizon 2020 research and innovation programme (Marie Sklodowska-Curie Global Fellowship - CANSAS, GA845065). This work was also supported in part by National Institutes of Health grants R01AG047589 to J.A.H.

## Author contributions

A.G. conceived the project and supervised research. LC designed and carried out the computational analyses, with input from MP and BB. JDW and JAH provided input on the use and interpretation of the voxel-scale anatomical connectivity data, and on the manuscript. AG and LC wrote the manuscript.

## Competing interests

The authors declare no competing interests.

## Materials & Correspondence

Correspondence and material request should be addressed to

## Supplementary Figures

**Figure S1.**
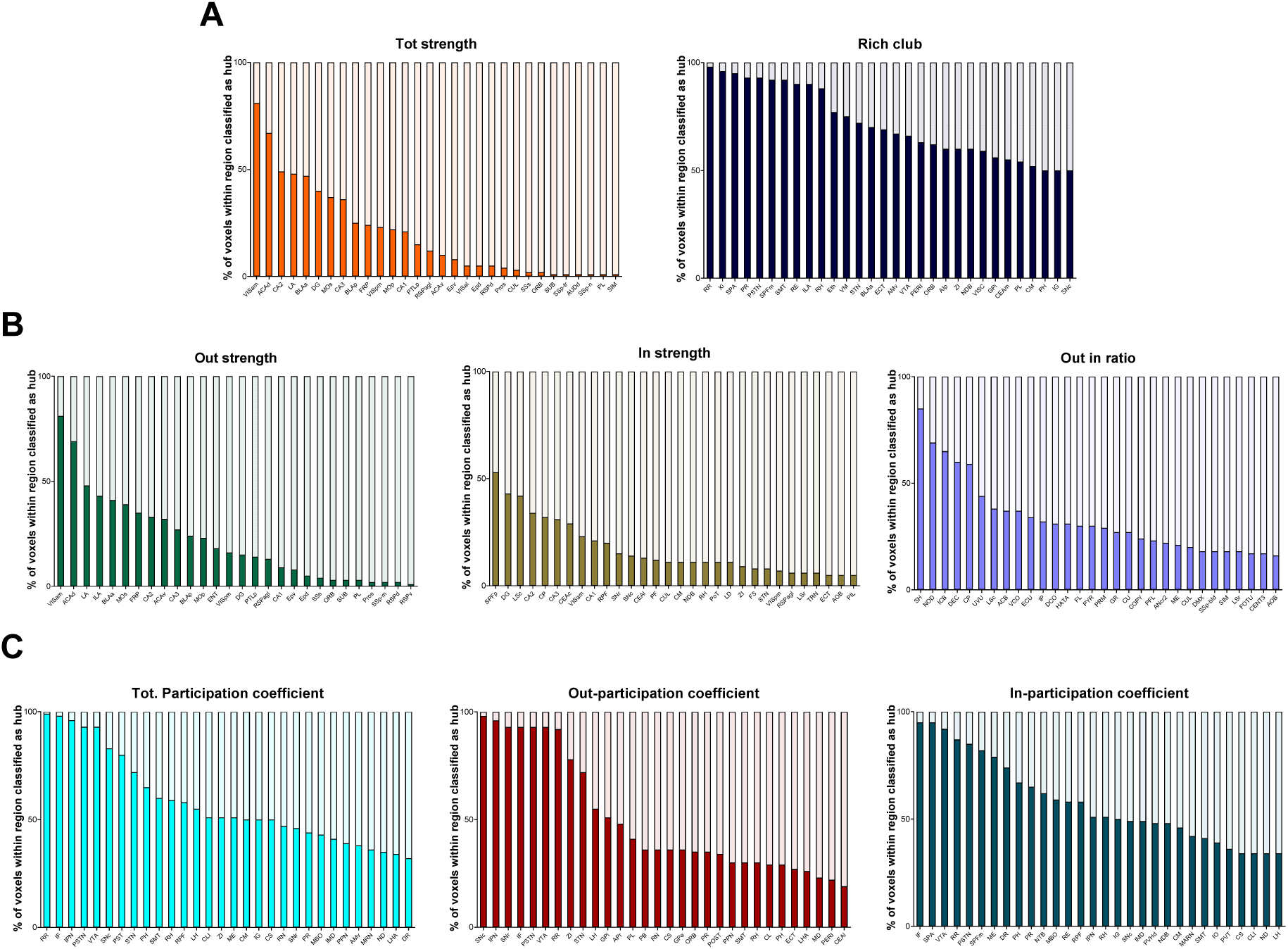
Regional quantification of hub‐like properties. (A ‐ C) For each measure of interest, we quantified the percentage of voxels classified as hub within their corresponding anatomical area. Anatomical regions were identified using the finest‐grained, volumetrically‐ matched parcellation of the mouse brain available (CCFv3, Knox et al., 2019). Values are expressed as % of the whole volume of the region of interest. The reported quantifications are here ranked in decreasing % of voxel exhibiting hub‐like properties within each area. Only the top 30 areas for each hub attribute were plotted representing as such the set of regions characterized by the lowest sub‐regional involvement for each metric.

**Figure S2.**
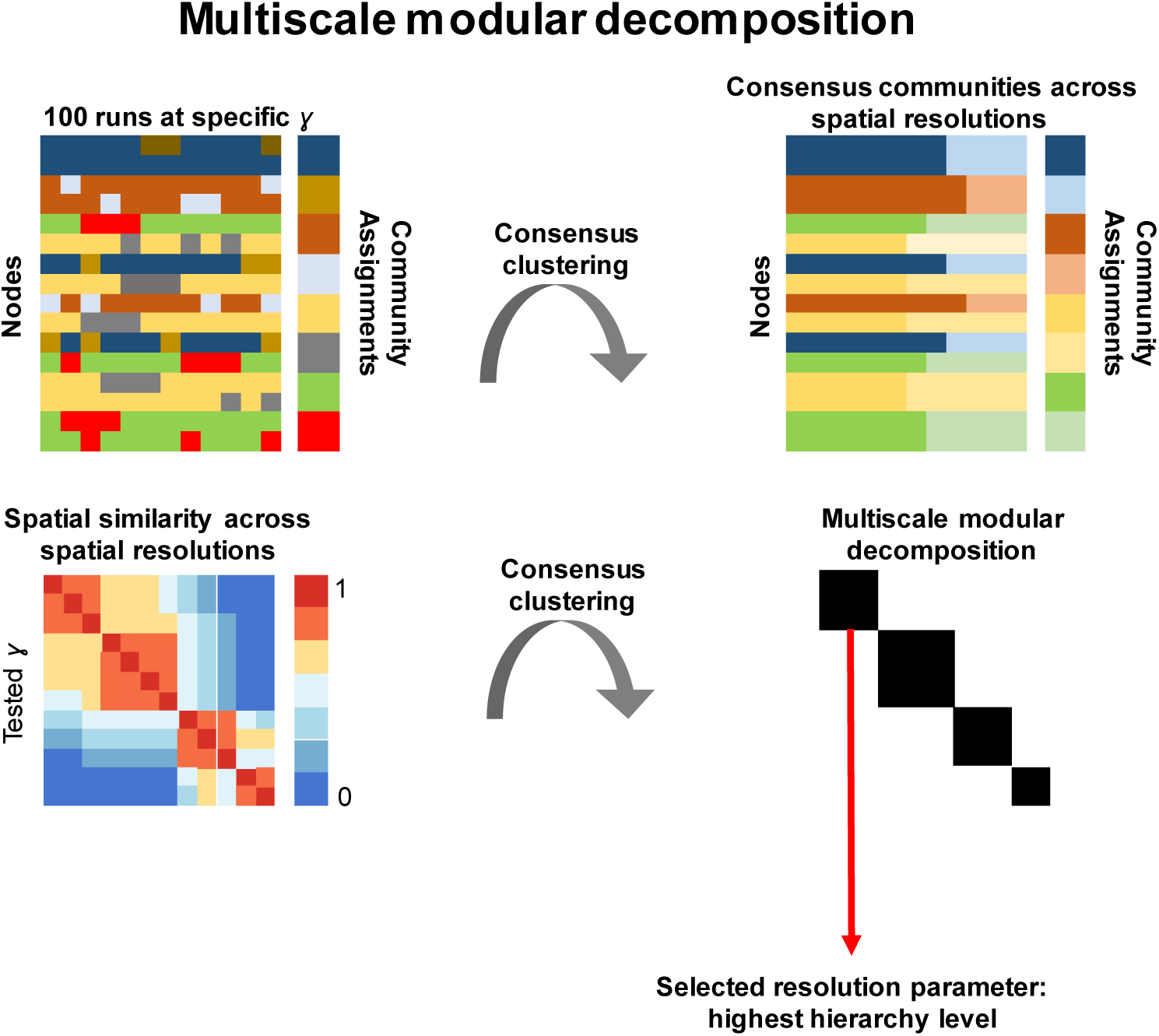
Multiscale modular decomposition of the structural connectome. We systematically varied the resolution parameter controlling the size of the modules, performing a consensus clustering and thus obtaining a representative community subdivision for each of the tested resolution setting as originally described by Rubinov et al., (2015), (top row). We then computed similarity between all the consensus partitions using the adjusted mutual information score. To avoid arbitrary thresholding of the similarity matrix, we again performed consensus clustering, obtaining at each step a representative hierarchical modular subdivision of the mouse structural connectome. We finally computed the agreement matrix, which encodes how many times any two hierarchical subdivisions of the connectome are grouped together (bottom row). We based all subsequent analyses on the partition obtained at the highest hierarchical level, testing its statistical significance against a set 1,000 rewired network with preserved degree and strength sequences, and where we additionally controlled for nodal wiring length, as assessed by Euclidean distance.

**Figure S3.**
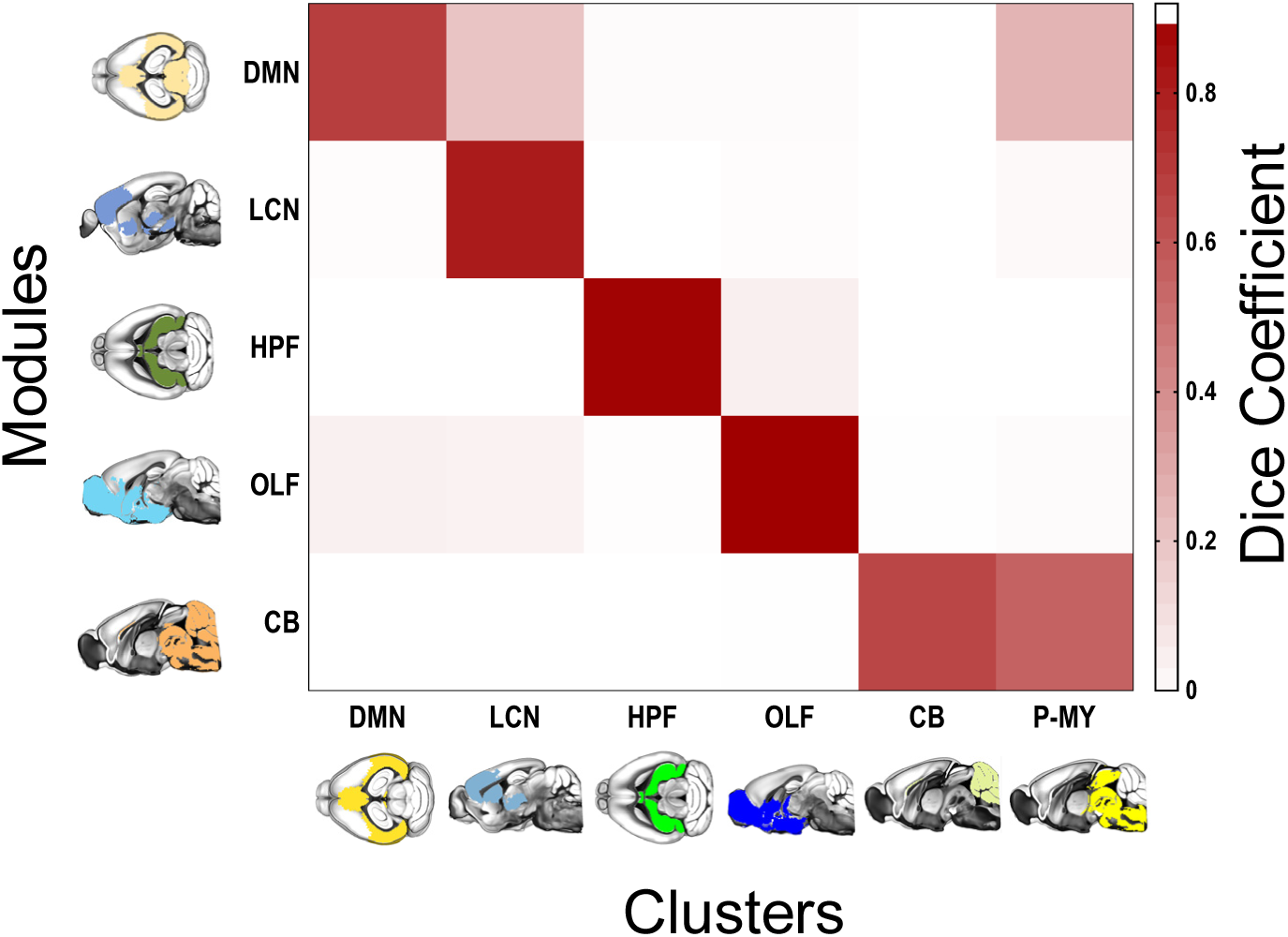
Agglomerative hierarchical clustering corroborates the multiscale modular decomposition approach. Quantification (Dice coefficient) of the spatial overlap between the modules obtained through the multiscale modular decomposition approach (left) and the most representative solution from an agglomerative hierarchical clustering (bottom). P‐MY: pons, medulla.

**Figure S4.**
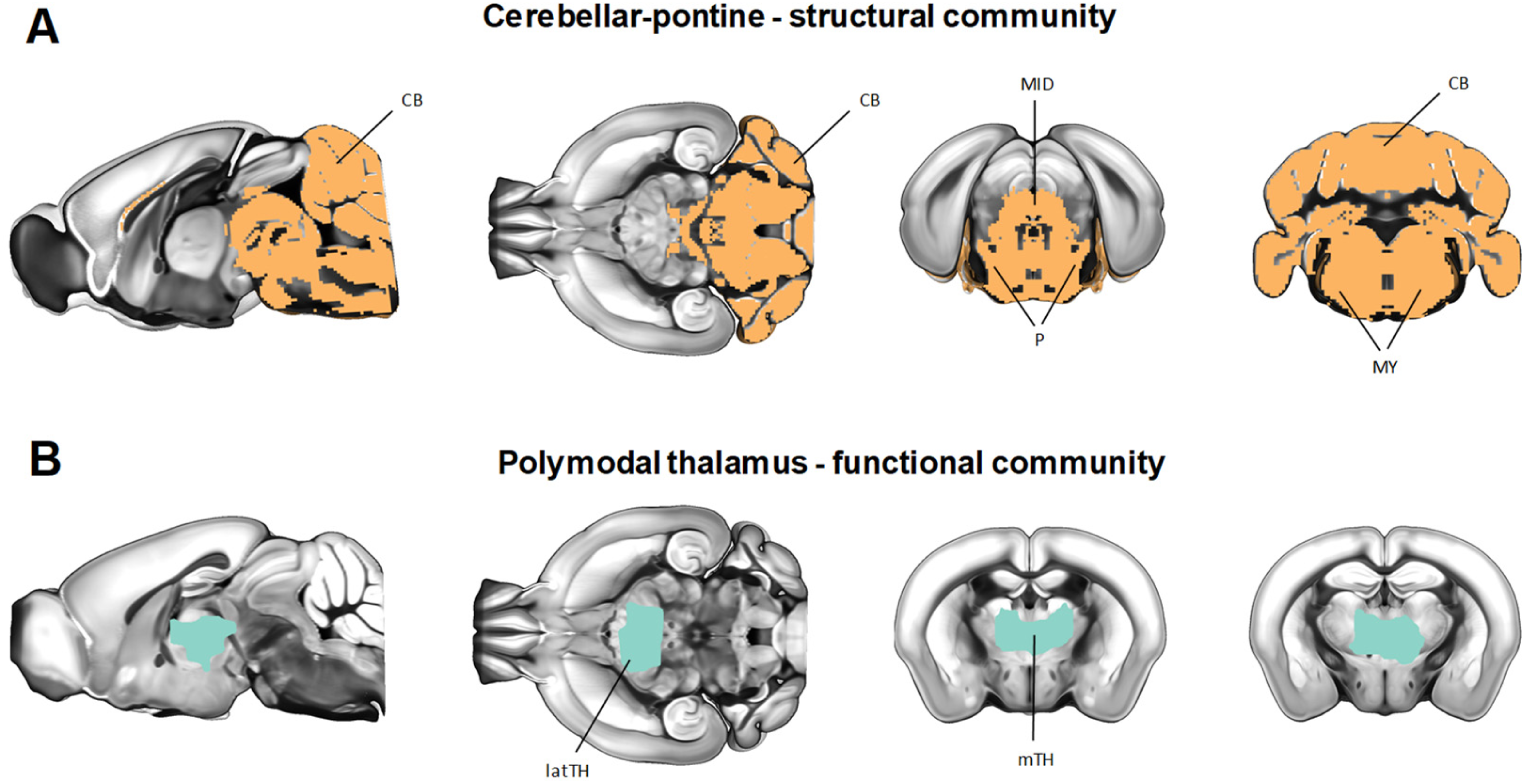
Non‐matching structural and functional communities. **(A)** Structural community spanning the cerebellum and pons (B) Thalamic functional community described in Liska et al. (2015)

**Figure S5.**
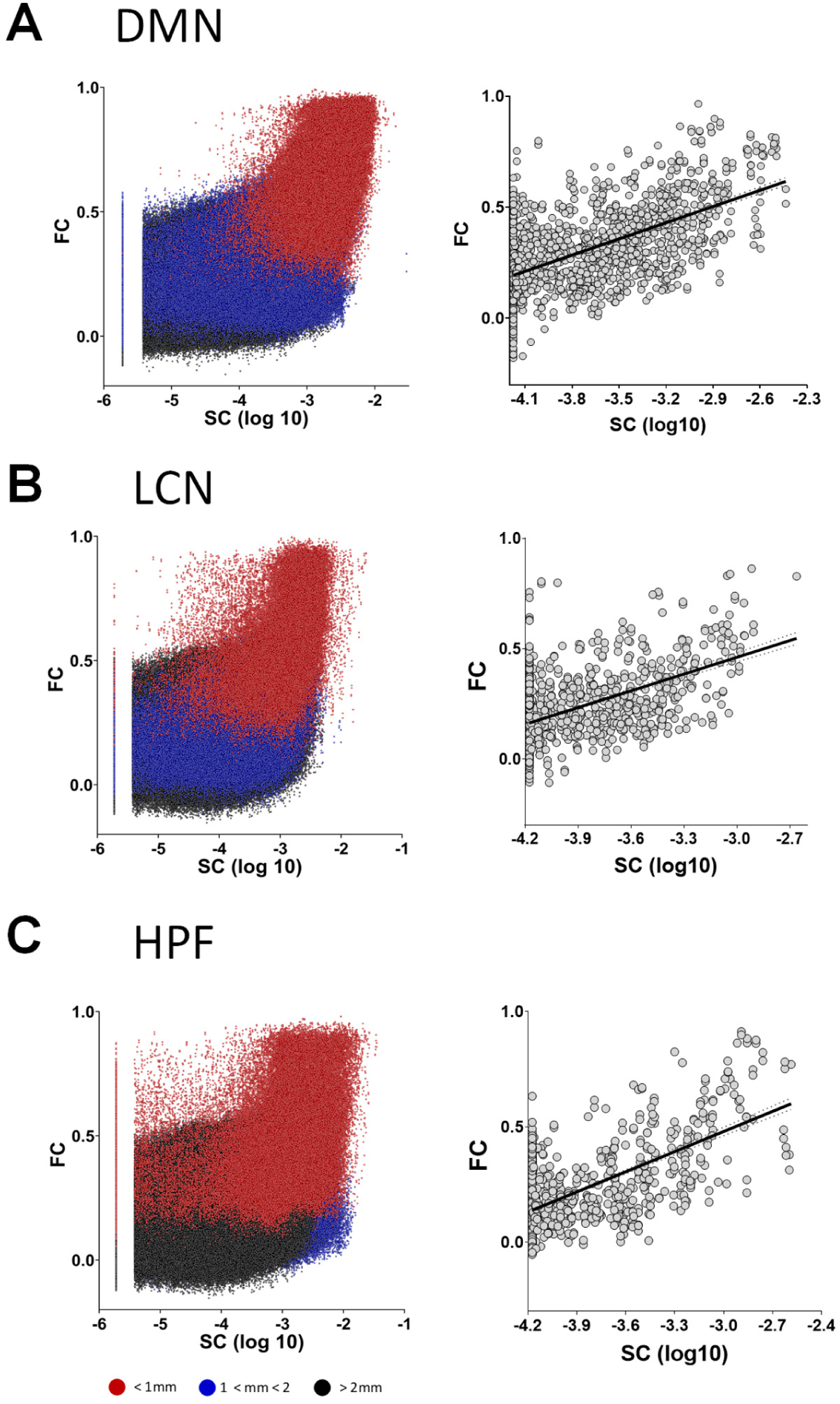
Correlation between structural (SC) and functional connectivity (FC) is distance dependent. (A) SC‐FC correlation for the Default Mode Network (DMN) module obtained by Liska et al., (2015). The correlation was computed at both the voxel level (left panel) and after aggregation in regions of interest (right panel) (B) SC‐FC correlation for the Latero cortical network (LCN) module obtained by Liska et al., (2015). The correlation was computed at both the voxel level (left panel) and after aggregation in regions of interest (right panel) (C) SC‐FC correlation for the hippocampal formation module (HPF) obtained by Liska et al., (2015). The correlation was computed at both the voxel level (left panel) and after aggregation in regions of interest (right panel). Red dots: distance < 1mm, Blue: distance >= 1mm and < 2mm, Black dots: distance >=2mm

**Figure S6.**
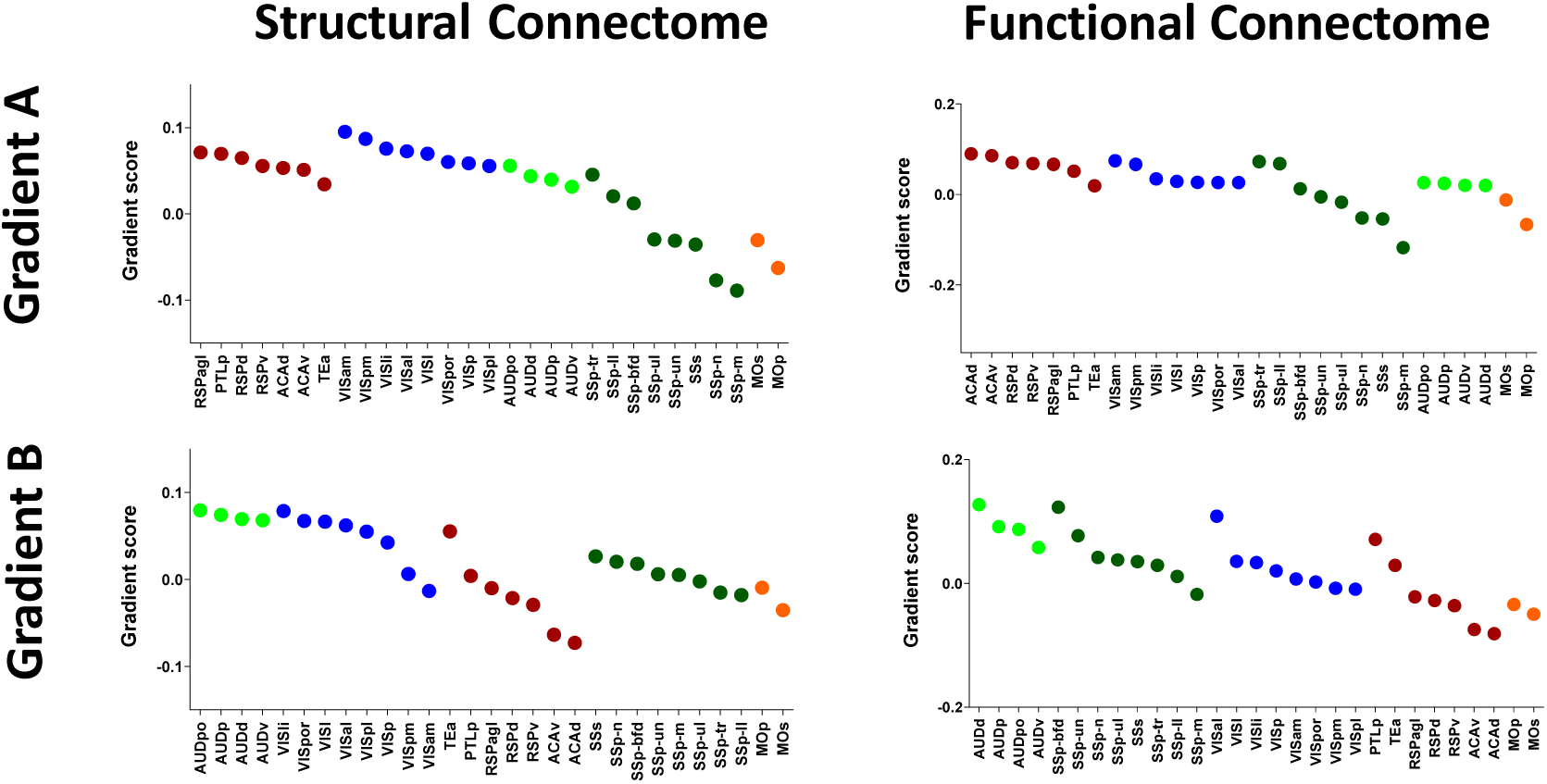
Regional quantification of gradient topography. A gradient participation score was quantified in each cortical region for both Gradient A (top row) and Gradient B (bottom row), both for the structural (Left column) and functional (Right column) connectomes. Anatomical abbreviations are defined in Supplementary Table I.

