## Supplementary material for "Network structure of the mouse brain connectome with voxel resolution": Table S1

| Abbreviation | Name |
|---|---|
| ACA | Anterior cingulate area |
| ACAd | Anterior cingulate area, dorsal part |
| ACAv | Anterior cingulate area, ventral part |
| Acb | Nucleus accumbens |
| AI | Agranular insular area |
| Amy | Amygdala |
| AUDd | Dorsal auditory area |
| AUDp | Primary auditory area |
| AUDpo | Posterior auditory area |
| AUDv | Ventral auditory area |
| CEREB | Cerebellum |
| CS | Superior central nucleus raphe |
| dHP | Dorsal hippocampus |
| DMN | Default Mode Network |
| DRN | Dorsal nucleus raphe |
| ENT | Entorhinal area |
| GP | Globus Pallidus |
| Ha | Habenula |
| Hy | Hypothalamus |
| IL | Infralimbic area |
| LC | Locus Coeruleus |
| LCN | Latero Cortical Network |
| LHb | lateral habenula |
| MD | Mediodorsal nucleus of thalamus |
| MOp | Primary motor area |
| MOs | Secondary motor area |
| OLF | Olfactory areas |
| ORB | Orbital area |
| PIR | Piriform area |
| PL | Prelimbic area |
| PPC | Posterior Parietal cortex |
| PTLp | Posterior parietal association areas |
| RE | Nucleus of Reuniens |
| RSP | Retrosplenial area |
| RSPagl | Retrosplenial area, lateral agranular part |
| RSPd | Retrosplenial area, dorsal part |
| RSPv | Retrosplenial area, ventral part |
| SEP | Septal complex |
| SN | Substantia Nigra |
| SSp | Primary somatosensory area |
| SSp-bfd | Primary somatosensory area, barrel field |
| SSp-ll | Primary somatosensory area, lower limb |
| SSp-m | Primary somatosensory area, mouth |
| SSp-n | Primary somatosensory area, nose |
| SSp-tr | Primary somatosensory area, trunk |
| SSp-ul | Primary somatosensory area, upper limb |

| | |
|---|---|
| SSp-un | Primary somatosensory area, unassigned |
| SSs | Supplemental somatosensory area |
| STR | Striatum |
| STRd | dorsal Striatum |
| STRv | ventral Striatum |
| TeA | Temporal association area |
| vHP | ventral Hippocampus |
| Vis | Visual area |
| VISal | Anterolateral visual area |
| VISam | Anteromedial visual area |
| VISl | Lateral visual area |
| VISli | Laterointermediate area |
| VISp | Primary visual area |
| VISpl | Posterolateral visual area |
| VISpm | posteromedial visual area |
| VISpor | Postrhinal area |
| VTA | Ventral tegmental area |
| ZI | Zona Incerta |
