## Supplementary material for "Network structure of the mouse brain connectome with voxel resolution": Table S2

| ID | ACRO | NAME |
|---|---|---|
| 184 | FRP | Frontal Pole |
| 985 | MOp | Primary motor area |
| 993 | MOs | Secondary motor area |
| 353 | SSp-n | Primary somatosensory area, nose |
| 329 | SSp-bfd | Primary somatosensory area, barrel field |
| 337 | SSp-ll | Primary somatosensory area, lower limb |
| 345 | SSp-m | Primary somatosensory area, mouth |
| 369 | SSp-ul | Primary somatosensory area, upper limb |
| 361 | SSp-tr | Primary somatosensory area, trunk |
| 182305689 | SSp-un | Primary somatosensory area, unassigned |
| 378 | SSs | Supplemental somatosensory area |
| 1057 | GU | Gustatory areas |
| 677 | VISC | Visceral area |
| 1011 | AUDd | Dorsal auditory area |
| 1002 | AUDp | Primary auditory area |
| 1027 | AUDpo | Posterior auditory area |
| 1018 | AUDv | Ventral auditory area |
| 402 | VISal | Anterolateral visual area |
| 394 | VISam | Anteromedial visual area |
| 409 | VISl | Lateral visual area |
| 385 | VISp | Primary visual area |
| 425 | VISpl | Posterolateral visual area |
| 533 | VISpm | posteromedial visual area |
| 312782574 | VISli | Laterointermediate area |
| 312782628 | VISpor | Postrhinal area |
| 39 | ACAd | Anterior cingulate area, dorsal part |
| 48 | ACAv | Anterior cingulate area, ventral part |
| 972 | PL | Prelimbic area |
| 44 | ILA | Infralimbic area |
| 714 | ORB | Orbital area |
| 104 | AId | Agranular insular area, dorsal part |
| 111 | AIp | Agranular insular area, posterior part |
| 119 | AIv | Agranular insular area, ventral part |
| 894 | RSPagl | Retrosplenial area, lateral agranular part |
| 879 | RSPd | Retrosplenial area, dorsal part |
| 886 | RSPv | Retrosplenial area, ventral part |
| 22 | PTLp | Posterior parietal association areas |
| 541 | TEa | Temporal association areas |
| 922 | PERI | Perirhinal area |
| 895 | ECT | Ectorhinal area |
| 507 | MOB | Main olfactory bulb |
| 151 | AOB | Accessory olfactory bulb |
| 159 | AON | Anterior olfactory nucleus |
| 589 | TT | Taenia tecta |
| 814 | DP | Dorsal peduncular area |
| 961 | PIR | Piriform area |
| 619 | NLOT | Nucleus of the lateral olfactory tract |
| 639 | COAa | Cortical amygdalar area, anterior part |
| 647 | COAp | Cortical amygdalar area, posterior part |
| 788 | PAA | Piriform-amygdalar area |
| 566 | TR | Postpiriform transition area |
| 382 | CA1 | Field CA1 |
| 423 | CA2 | Field CA2 |
| 463 | CA3 | Field CA3 |

| | | |
|---|---|---|
| 726 | DG | Dentate gyrus |
| 982 | FC | Fasciola cinerea |
| 19 | IG | Induseum griseum |
| 909 | ENT | Entorhinal area |
| 843 | PAR | Parasubiculum |
| 1037 | POST | Postsubiculum |
| 1084 | PRE | Presubiculum |
| 502 | SUB | Subiculum |
| 484682470 | Pros | Prosubiculum |
| 589508447 | HATA | Hippocampo-amygdalar transition area |
| 484682508 | APr | Area prostriata |
| 583 | CLA | Claustrum |
| 952 | Epd | Endopiriform nucleus, dorsal part |
| 966 | Epv | Endopiriform nucleus, ventral part |
| 131 | LA | Lateral amygdalar nucleus |
| 303 | BLAa | Basolateral amygdalar nucleus, anterior part |
| 311 | BLAp | Basolateral amygdalar nucleus, posterior part |
| 451 | BLAv | Basolateral amygdalar nucleus, ventral part |
| 327 | BMAa | Basomedial amygdalar nucleus, anterior part |
| 334 | BMAp | Basomedial amygdalar nucleus, posterior part |
| 780 | PA | Posterior amygdalar nucleus |
| 672 | CP | Caudoputamen |
| 56 | ACB | Nucleus accumbens |
| 998 | FS | Fundus of striatum |
| 754 | OT | Olfactory tubercle |
| 250 | LSc | Lateral septal nucleus, caudal (caudodorsal) part |
| 258 | LSr | Lateral septal nucleus, rostral (rostroventral) part |
| 266 | LSv | Lateral septal nucleus, ventral part |
| 310 | SF | Septofimbrial nucleus |
| 333 | SH | Septohippocampal nucleus |
| 23 | AAA | Anterior amygdalar area |
| 292 | BA | Bed nucleus of the accessory olfactory tract |
| 544 | CEAc | Central amygdalar nucleus, capsular part |
| 551 | CEAl | Central amygdalar nucleus, lateral part |
| 559 | CEAm | Central amygdalar nucleus, medial part |
| 1105 | IA | Intercalated amygdalar nucleus |
| 403 | MEA | Medial amygdalar nucleus |
| 1022 | GPe | Globus pallidus, external segment |
| 1031 | GPi | Globus pallidus, internal segment |
| 342 | SI | Substantia innominata |
| 298 | MA | Magnocellular nucleus |
| 564 | MS | Medial septal nucleus |
| 596 | NDB | Diagonal band nucleus |
| 351 | BST | Bed nuclei of the stria terminalis |
| 287 | BAC | Bed nucleus of the anterior commissure |
| 629 | VAL | Ventral anterior-lateral complex of the thalamus |
| 685 | VM | Ventral medial nucleus of the thalamus |
| 709 | VP | Ventral posterior complex of the thalamus |
| 563807435 | PoT | Posterior triangular thalamic nucleus |
| 414 | SPFm | Subparafascicular nucleus, magnocellular part |
| 422 | SPFp | Subparafascicular nucleus, parvicellular part |
| 609 | SPA | Subparafascicular area |
| 1044 | PP | Peripeduncular nucleus |
| 1072 | MGd | Medial geniculate complex, dorsal part |
| 1079 | MGv | Medial geniculate complex, ventral part |

| | | |
|---|---|---|
| 1088 | MGm | Medial geniculate complex, medial part |
| 496345664 | LGd-sh | Dorsal part of the lateral geniculate complex, shell |
| 496345668 | LGd-co | Dorsal part of the lateral geniculate complex, core |
| 496345672 | LGd-ip | Dorsal part of the lateral geniculate complex, ipsilateral zone |
| 218 | LP | Lateral posterior nucleus of the thalamus |
| 1020 | PO | Posterior complex of the thalamus |
| 1029 | POL | Posterior limiting nucleus of the thalamus |
| 325 | SGN | Suprageniculate nucleus |
| 560581551 | Eth | Ethmoid nucleus of the thalamus |
| 255 | AV | Anteroventral nucleus of thalamus |
| 1096 | AMd | Anteromedial nucleus, dorsal part |
| 1104 | AMv | Anteromedial nucleus,ventral part |
| 64 | AD | Anterodorsal nucleus |
| 1120 | IAM | Interanteromedial nucleus of the thalamus |
| 1113 | IAD | Interanterodorsal nucleus of the thalamus |
| 155 | LD | Lateral dorsal nucleus of thalamus |
| 59 | IMD | Intermediodorsal nucleus of the thalamus |
| 362 | MD | Mediodorsal nucleus of thalamus |
| 366 | SMT | Submedial nucleus of the thalamus |
| 1077 | PR | Perireunensis nucleus |
| 149 | PVT | Paraventricular nucleus of the thalamus |
| 15 | PT | Parataenial nucleus |
| 181 | RE | Nucleus of reuniens |
| 560581559 | Xi | Xiphoid thalamic nucleus |
| 189 | RH | Rhomboid nucleus |
| 599 | CM | Central medial nucleus of the thalamus |
| 907 | PCN | Paracentral nucleus |
| 575 | CL | Central lateral nucleus of the thalamus |
| 930 | PF | Parafascicular nucleus |
| 560581563 | PIL | Posterior intralaminar thalamic nucleus |
| 262 | RT | Reticular nucleus of the thalamus |
| 27 | IGL | Intergeniculate leaflet of the lateral geniculate complex |
| 563807439 | IntG | Intermediate geniculate nucleus |
| 178 | Lgv | Ventral part of the lateral geniculate complex |
| 321 | SubG | Subgeniculate nucleus |
| 483 | MH | Medial habenula |
| 186 | LH | Lateral habenula |
| 390 | SO | Supraoptic nucleus |
| 332 | ASO | Accessory supraoptic group |
| 38 | PVH | Paraventricular hypothalamic nucleus |
| 30 | Pva | Periventricular hypothalamic nucleus, anterior part |
| 118 | Pvi | Periventricular hypothalamic nucleus, intermediate part |
| 223 | ARH | Arcuate hypothalamic nucleus |
| 72 | ADP | Anterodorsal preoptic nucleus |
| 263 | AVP | Anteroventral preoptic nucleus |
| 272 | AVPV | Anteroventral periventricular nucleus |
| 830 | DMH | Dorsomedial nucleus of the hypothalamus |
| 452 | MEPO | Median preoptic nucleus |
| 523 | MPO | Medial preoptic area |
| 763 | OV | Vascular organ of the lamina terminalis |
| 914 | PD | Posterodorsal preoptic nucleus |
| 1109 | PS | Parastrial nucleus |
| 126 | PVp | Periventricular hypothalamic nucleus, posterior part |
| 133 | PVpo | Periventricular hypothalamic nucleus, preoptic part |
| 347 | SBPV | Subparaventricular zone |

| | | |
|---|---|---|
| 286 | SCH | Suprachiasmatic nucleus |
| 338 | SFO | Subfornical organ |
| 576073699 | VMPO | Ventromedial preoptic nucleus |
| 689 | VLPO | Ventrolateral preoptic nucleus |
| 88 | AHN | Anterior hypothalamic nucleus |
| 331 | MBO | Mammillary body |
| 515 | MPN | Medial preoptic nucleus |
| 980 | PMd | Dorsal premammillary nucleus |
| 1004 | Pmv | Ventral premammillary nucleus |
| 63 | PVHd | Paraventricular hypothalamic nucleus, descending division |
| 693 | VMH | Ventromedial hypothalamic nucleus |
| 946 | PH | Posterior hypothalamic nucleus |
| 194 | LHA | Lateral hypothalamic area |
| 226 | LPO | Lateral preoptic area |
| 356 | PST | Preparasubthalamic nucleus |
| 364 | PSTN | Parasubthalamic nucleus |
| 576073704 | PeF | Perifornical nucleus |
| 173 | RCH | Retrochiasmatic area |
| 470 | STN | Subthalamic nucleus |
| 614 | TU | Tuberal nucleus |
| 797 | ZI | Zona incerta |
| 10671 | ME | Median eminence |
| 302 | SCs | Superior colliculus, sensory related |
| 811 | ICc | Inferior colliculus, central nucleus |
| 820 | Icd | Inferior colliculus,dorsal nucleus |
| 828 | Ice | Inferior colliculus, external |
| 580 | NB | Nucleus of the brachium of the inferior colliculus |
| 271 | SAG | Nucleus sagulum |
| 874 | PBG | Parabigeminal nucleus |
| 460 | MEV | Midbrain trigeminal nucleus |
| 599626923 | SCO | Subcommissural organ |
| 381 | SNr | Substantia nigra, reticular part |
| 749 | VTA | Ventral tegmental area |
| 607344830 | PN | Paranigral nucleus |
| 246 | RR | Midbrain reticular nucleus, retrorubral area |
| 128 | MRN | Midbrain reticular nucleus |
| 294 | SCm | Superior colliculus, motor related |
| 50 | PRC | Precommissural nucleus |
| 67 | INC | Interstitial nucleus of Cajal |
| 587 | ND | Nucleus of Darkschewitsch |
| 614454277 | SU3 | Supraoculomotor periaqueductal gray |
| 215 | APN | Anterior pretectal nucleus |
| 531 | MPT | Medial pretectal area |
| 628 | NOT | Nucleus of the optic tract |
| 634 | NPC | Nucleus of the posterior commissure |
| 706 | OP | Olivary pretectal nucleus |
| 1061 | PPT | Posterior pretectal nucleus |
| 549009203 | RPF | Retroparafascicular nucleus |
| 616 | CUN | Cuneiform nucleus |
| 214 | RN | Red nucleus |
| 35 | III | Oculomotor nucleus |
| 549009211 | MA3 | Medial accesory oculomotor nucleus |
| 975 | EW | Edinger-Westphal nucleus |
| 115 | IV | Trochlear nucleus |
| 757 | VTN | Ventral tegmental nucleus |

| | | |
|---|---|---|
| 231 | AT | Anterior tegmental nucleus |
| 66 | LT | Lateral terminal nucleus of the accessory optic tract |
| 75 | DT | Dorsal terminal nucleus of the accessory optic tract |
| 58 | MT | Medial terminal nucleus of the accessory optic tract |
| 374 | SNc | Substantia nigra, compact part |
| 1052 | PPN | Pedunculopontine nucleus |
| 12 | IF | Interfascicular nucleus raphe |
| 100 | IPN | Interpeduncular nucleus |
| 197 | RL | Rostral linear nucleus raphe |
| 591 | CLI | Central linear nucleus raphe |
| 872 | DR | Dorsal nucleus raphe |
| 612 | NLL | Nucleus of the lateral lemniscus |
| 7 | PSV | Principal sensory nucleus of the trigeminal |
| 867 | PB | Parabrachial nucleus |
| 122 | POR | Superior olivary complex, periolivary region |
| 105 | SOCm | Superior olivary complex, medial part |
| 114 | SOCl | Superior olivary complex, lateral part |
| 280 | B | Barrington's nucleus |
| 880 | DTN | Dorsal tegmental nucleus |
| 599626927 | PDTg | Posterodorsal tegmental nucleus |
| 898 | PCG | Pontine central gray |
| 1093 | PRNc | Pontine reticular nucleus, caudal part |
| 318 | SG | Supragenual nucleus |
| 534 | SUT | Supratrigeminal nucleus |
| 574 | TRN | Tegmental reticular nucleus |
| 621 | V | Motor nucleus of trigeminal |
| 549009215 | P5 | Peritrigeminal zone |
| 549009219 | Acs5 | Accessory trigeminal nucleus |
| 549009223 | PC5 | Parvicellular motor 5 nucleus |
| 549009227 | I5 | Intertrigeminal nucleus |
| 679 | CS | Superior central nucleus raphe |
| 147 | LC | Locus ceruleus |
| 162 | LDT | Laterodorsal tegmental nucleus |
| 604 | NI | Nucleus incertus |
| 146 | PRNr | Pontine reticular nucleus |
| 238 | RPO | Nucleus raphe pontis |
| 350 | SLC | Subceruleus nucleus |
| 358 | SLD | Sublaterodorsal nucleus |
| 207 | AP | Area Postrema |
| 96 | DCO | Dorsal cochlear nucleus |
| 101 | VCO | Ventral cochlear nucleus |
| 711 | CU | Cuneate nucleus |
| 1039 | GR | Gracile nucleus |
| 903 | ECU | External cuneate nucleus |
| 642 | NTB | Nucleus of the trapezoid body |
| 651 | NTS | Nucleus of the solitary tract |
| 429 | SPVc | Spinal nucleus of the trigeminal, caudal part |
| 437 | SPVI | Spinal nucleus of the trigeminal, interpolar part |
| 445 | SPVO | Spinal nucleus of the trigeminal, oral part |
| 589508451 | Pa5 | Paratrigeminal nucleus |
| 653 | VI | Abducens nucleus |
| 661 | VII | Facial motor nucleus |
| 576 | ACVII | Accessory facial motor nucleus |
| 939 | AMBd | Nucleus ambiguus, dorsal division |
| 143 | AMBv | Nucleus ambiguus, ventral division |

| | | |
|---|---|---|
| 839 | DMX | Dorsal motor nucleus of the vagus nerve |
| 1048 | GRN | Gigantocellular reticular nucleus |
| 372 | ICB | Infracerebellar nucleus |
| 83 | IO | Inferior olivary complex |
| 136 | IRN | Intermediate reticular nucleus |
| 106 | ISN | Inferior salivatory nucleus |
| 203 | LIN | Linear nucleus of the medulla |
| 955 | LRNm | Lateral reticular nucleus, magnocellular part |
| 963 | LRNp | Lateral reticular nucleus, parvicellular part |
| 307 | MARN | Magnocellular reticular nucleus |
| 1098 | MDRNd | Medullary reticular nucleus, dorsal part |
| 1107 | MDRNv | Medullary reticular nucleus, ventral part |
| 852 | PARN | Parvicellular reticular nucleus |
| 859 | PAS | Parasolitary nucleus |
| 970 | PGRNd | Paragigantocellular reticular nucleus, dorsal part |
| 978 | PGRNl | Paragigantocellular reticular nucleus, lateral part |
| 177 | NR | Nucleus of Roller |
| 169 | PRP | Nucleus prepositus |
| 1069 | PPY | Parapyramidal nucleus |
| 209 | LAV | Lateral vestibular nucleus |
| 202 | MV | Medial vestibular nucleus |
| 225 | SPIV | Spinal vestibular nucleus |
| 217 | SUV | Superior vestibular nucleus |
| 765 | x | Nucleus x |
| 773 | XII | Hypoglossal nucleus |
| 781 | y | Nucleus y |
| 206 | RM | Nucleus raphe magnus |
| 230 | RPA | Nucleus raphe pallidus |
| 222 | RO | Nucleus raphe obscurus |
| 912 | LING | Lingula (I) |
| 976 | CENT2 | Lobule II |
| 984 | CENT3 | Lobule III |
| 928 | CUL | Culmen |
| 936 | DEC | Declive (VI) |
| 944 | FOTU | Folium-tuber vermis (VII) |
| 951 | PYR | Pyramus (VIII) |
| 957 | UVU | Uvula (IX) |
| 968 | NOD | Nodulus (X) |
| 1007 | SIM | Simple lobule |
| 1056 | ANcr1 | Crus 1 |
| 1064 | ANcr2 | Crus 2 |
| 1025 | PRM | Paramedian lobule |
| 1033 | COPY | Copula pyramidis |
| 1041 | PFL | Paraflocculus |
| 1049 | FL | Flocculus |
| 989 | FN | Fastigial nucleus |
| 91 | IP | Interposed nucleus |
| 846 | DN | Dentate nucleus |
| 589508455 | VeCB | Vestibulocerebellar nucleus |

| Macro | # voxels bilateral | # voxel right | # voxel left |
|---|---|---|---|
| Isocortex | 966 | 483 | 483 |
| Isocortex | 11376 | 5718 | 5658 |
| Isocortex | 13096 | 6552 | 6544 |
| Isocortex | 3032 | 1520 | 1512 |
| Isocortex | 6281 | 3137 | 3144 |
| Isocortex | 2361 | 1178 | 1183 |
| Isocortex | 6224 | 3115 | 3109 |
| Isocortex | 3764 | 1878 | 1886 |
| Isocortex | 1399 | 703 | 696 |
| Isocortex | 1263 | 633 | 630 |
| Isocortex | 8993 | 4504 | 4489 |
| Isocortex | 1760 | 883 | 877 |
| Isocortex | 2370 | 1184 | 1186 |
| Isocortex | 1213 | 609 | 604 |
| Isocortex | 2152 | 1079 | 1073 |
| Isocortex | 598 | 300 | 298 |
| Isocortex | 1807 | 904 | 903 |
| Isocortex | 768 | 391 | 377 |
| Isocortex | 775 | 386 | 389 |
| Isocortex | 1237 | 618 | 619 |
| Isocortex | 7113 | 3565 | 3548 |
| Isocortex | 792 | 399 | 393 |
| Isocortex | 1043 | 520 | 523 |
| Isocortex | 492 | 243 | 249 |
| Isocortex | 1269 | 638 | 631 |
| Isocortex | 3114 | 1479 | 1635 |
| Isocortex | 2387 | 1128 | 1259 |
| Isocortex | 2433 | 1159 | 1274 |
| Isocortex | 849 | 403 | 446 |
| Isocortex | 5886 | 2907 | 2979 |
| Isocortex | 3726 | 1867 | 1859 |
| Isocortex | 2429 | 1211 | 1218 |
| Isocortex | 1737 | 873 | 864 |
| Isocortex | 2308 | 1162 | 1146 |
| Isocortex | 3816 | 1899 | 1917 |
| Isocortex | 4331 | 2090 | 2241 |
| Isocortex | 2454 | 1232 | 1222 |
| Isocortex | 3106 | 1549 | 1557 |
| Isocortex | 797 | 398 | 399 |
| Isocortex | 1728 | 870 | 858 |
| Olfactory Areas | 16406 | 8218 | 8188 |
| Olfactory Areas | 650 | 325 | 325 |
| Olfactory Areas | 4880 | 2437 | 2443 |
| Olfactory Areas | 1431 | 690 | 741 |
| Olfactory Areas | 482 | 232 | 250 |
| Olfactory Areas | 11591 | 5793 | 5798 |
| Olfactory Areas | 311 | 153 | 158 |
| Olfactory Areas | 763 | 387 | 376 |
| Olfactory Areas | 2495 | 1251 | 1244 |
| Olfactory Areas | 1235 | 619 | 616 |
| Olfactory Areas | 1323 | 658 | 665 |
| Hippocampal formation | 10278 | 5145 | 5133 |
| Hippocampal formation | 451 | 226 | 225 |
| Hippocampal formation | 6289 | 3143 | 3146 |

| | | | |
|---|---|---|---|
| Hippocampal formation | 6571 | 3275 | 3296 |
| Hippocampal formation | 57 | 29 | 28 |
| Hippocampal formation | 108 | 40 | 68 |
| Hippocampal formation | 11476 | 5741 | 5735 |
| Hippocampal formation | 930 | 467 | 463 |
| Hippocampal formation | 1074 | 535 | 539 |
| Hippocampal formation | 906 | 454 | 452 |
| Hippocampal formation | 2146 | 1070 | 1076 |
| Hippocampal formation | 1185 | 594 | 591 |
| Hippocampal formation | 420 | 213 | 207 |
| Hippocampal formation | 361 | 179 | 182 |
| Cortical subplate | 545 | 271 | 274 |
| Cortical subplate | 1796 | 899 | 897 |
| Cortical subplate | 961 | 476 | 485 |
| Cortical subplate | 843 | 424 | 419 |
| Cortical subplate | 764 | 379 | 385 |
| Cortical subplate | 710 | 361 | 349 |
| Cortical subplate | 414 | 210 | 204 |
| Cortical subplate | 777 | 387 | 390 |
| Cortical subplate | 708 | 352 | 356 |
| Cortical subplate | 966 | 481 | 485 |
| Striatum | 26040 | 13031 | 13009 |
| Striatum | 4446 | 2224 | 2222 |
| Striatum | 424 | 212 | 212 |
| Striatum | 3829 | 1913 | 1916 |
| Striatum | 572 | 288 | 284 |
| Striatum | 1896 | 939 | 957 |
| Striatum | 601 | 300 | 301 |
| Striatum | 482 | 215 | 267 |
| Striatum | 33 | 16 | 17 |
| Striatum | 504 | 250 | 254 |
| Striatum | 25 | 12 | 13 |
| Striatum | 308 | 155 | 153 |
| Striatum | 267 | 130 | 137 |
| Striatum | 750 | 378 | 372 |
| Striatum | 179 | 89 | 90 |
| Striatum | 2024 | 1014 | 1010 |
| Pallidum | 1560 | 784 | 776 |
| Pallidum | 427 | 214 | 213 |
| Pallidum | 3000 | 1489 | 1511 |
| Pallidum | 367 | 185 | 182 |
| Pallidum | 405 | 147 | 258 |
| Pallidum | 723 | 333 | 390 |
| Pallidum | 1341 | 672 | 669 |
| Pallidum | 8 | 4 | 4 |
| Thalamus | 801 | 405 | 396 |
| Thalamus | 946 | 469 | 477 |
| Thalamus | 2852 | 1423 | 1429 |
| Thalamus | 281 | 141 | 140 |
| Thalamus | 66 | 33 | 33 |
| Thalamus | 144 | 72 | 72 |
| Thalamus | 139 | 57 | 82 |
| Thalamus | 58 | 28 | 30 |
| Thalamus | 166 | 84 | 82 |
| Thalamus | 276 | 138 | 138 |

| | | | |
|---|---|---|---|
| Thalamus | 255 | 127 | 128 |
| Thalamus | 205 | 103 | 102 |
| Thalamus | 420 | 209 | 211 |
| Thalamus | 88 | 44 | 44 |
| Thalamus | 1221 | 609 | 612 |
| Thalamus | 1268 | 630 | 638 |
| Thalamus | 204 | 103 | 101 |
| Thalamus | 184 | 93 | 91 |
| Thalamus | 240 | 117 | 123 |
| Thalamus | 415 | 208 | 207 |
| Thalamus | 247 | 124 | 123 |
| Thalamus | 172 | 86 | 86 |
| Thalamus | 164 | 83 | 81 |
| Thalamus | 47 | 20 | 27 |
| Thalamus | 120 | 59 | 61 |
| Thalamus | 1006 | 503 | 503 |
| Thalamus | 186 | 74 | 112 |
| Thalamus | 1373 | 678 | 695 |
| Thalamus | 297 | 150 | 147 |
| Thalamus | 150 | 75 | 75 |
| Thalamus | 444 | 178 | 266 |
| Thalamus | 230 | 113 | 117 |
| Thalamus | 436 | 203 | 233 |
| Thalamus | 74 | 14 | 60 |
| Thalamus | 93 | 38 | 55 |
| Thalamus | 256 | 108 | 148 |
| Thalamus | 231 | 118 | 113 |
| Thalamus | 351 | 180 | 171 |
| Thalamus | 507 | 258 | 249 |
| Thalamus | 168 | 84 | 84 |
| Thalamus | 1432 | 718 | 714 |
| Thalamus | 67 | 34 | 33 |
| Thalamus | 22 | 11 | 11 |
| Thalamus | 408 | 205 | 203 |
| Thalamus | 27 | 13 | 14 |
| Thalamus | 315 | 157 | 158 |
| Thalamus | 343 | 169 | 174 |
| Hypothalamus | 40 | 21 | 19 |
| Hypothalamus | 3 | 2 | 1 |
| Hypothalamus | 179 | 89 | 90 |
| Hypothalamus | 30 | 28 | 2 |
| Hypothalamus | 151 | 118 | 33 |
| Hypothalamus | 277 | 134 | 143 |
| Hypothalamus | 91 | 46 | 45 |
| Hypothalamus | 89 | 47 | 42 |
| Hypothalamus | 184 | 89 | 95 |
| Hypothalamus | 386 | 177 | 209 |
| Hypothalamus | 35 | 2 | 33 |
| Hypothalamus | 566 | 282 | 284 |
| Hypothalamus | 6 | 0 | 6 |
| Hypothalamus | 10 | 5 | 5 |
| Hypothalamus | 102 | 50 | 52 |
| Hypothalamus | 117 | 56 | 61 |
| Hypothalamus | 112 | 55 | 57 |
| Hypothalamus | 120 | 47 | 73 |

| | | | |
|---|---|---|---|
| Hypothalamus | 69 | 35 | 34 |
| Hypothalamus | 22 | 7 | 15 |
| Hypothalamus | 44 | 22 | 22 |
| Hypothalamus | 63 | 30 | 33 |
| Hypothalamus | 723 | 363 | 360 |
| Hypothalamus | 1017 | 478 | 539 |
| Hypothalamus | 401 | 191 | 210 |
| Hypothalamus | 138 | 70 | 68 |
| Hypothalamus | 194 | 97 | 97 |
| Hypothalamus | 126 | 62 | 64 |
| Hypothalamus | 543 | 270 | 273 |
| Hypothalamus | 701 | 338 | 363 |
| Hypothalamus | 2117 | 1062 | 1055 |
| Hypothalamus | 511 | 257 | 254 |
| Hypothalamus | 15 | 8 | 7 |
| Hypothalamus | 161 | 79 | 82 |
| Hypothalamus | 211 | 106 | 105 |
| Hypothalamus | 137 | 69 | 68 |
| Hypothalamus | 200 | 100 | 100 |
| Hypothalamus | 530 | 268 | 262 |
| Hypothalamus | 1815 | 910 | 905 |
| Hypothalamus | 85 | 34 | 51 |
| Midbrain | 2124 | 1042 | 1082 |
| Midbrain | 1120 | 558 | 562 |
| Midbrain | 1311 | 651 | 660 |
| Midbrain | 2012 | 1010 | 1002 |
| Midbrain | 89 | 46 | 43 |
| Midbrain | 107 | 52 | 55 |
| Midbrain | 49 | 25 | 24 |
| Midbrain | 13 | 7 | 6 |
| Midbrain | 13 | 2 | 11 |
| Midbrain | 1564 | 787 | 777 |
| Midbrain | 427 | 211 | 216 |
| Midbrain | 23 | 11 | 12 |
| Midbrain | 131 | 66 | 65 |
| Midbrain | 5174 | 2588 | 2586 |
| Midbrain | 5662 | 2789 | 2873 |
| Midbrain | 174 | 87 | 87 |
| Midbrain | 80 | 40 | 40 |
| Midbrain | 97 | 50 | 47 |
| Midbrain | 42 | 21 | 21 |
| Midbrain | 1277 | 639 | 638 |
| Midbrain | 47 | 23 | 24 |
| Midbrain | 214 | 108 | 106 |
| Midbrain | 286 | 144 | 142 |
| Midbrain | 60 | 30 | 30 |
| Midbrain | 143 | 72 | 71 |
| Midbrain | 64 | 31 | 33 |
| Midbrain | 564 | 284 | 280 |
| Midbrain | 792 | 396 | 396 |
| Midbrain | 32 | 17 | 15 |
| Midbrain | 20 | 10 | 10 |
| Midbrain | 19 | 0 | 19 |
| Midbrain | 6 | 3 | 3 |
| Midbrain | 34 | 17 | 17 |

| | | | |
|---|---|---|---|
| Midbrain | 43 | 21 | 22 |
| Midbrain | 17 | 9 | 8 |
| Midbrain | 12 | 6 | 6 |
| Midbrain | 51 | 25 | 26 |
| Midbrain | 203 | 101 | 102 |
| Midbrain | 888 | 442 | 446 |
| Midbrain | 84 | 34 | 50 |
| Midbrain | 348 | 147 | 201 |
| Midbrain | 57 | 17 | 40 |
| Midbrain | 82 | 35 | 47 |
| Midbrain | 151 | 37 | 114 |
| Pons | 739 | 369 | 370 |
| Pons | 1108 | 550 | 558 |
| Pons | 1136 | 565 | 570 |
| Pons | 372 | 185 | 187 |
| Pons | 198 | 98 | 100 |
| Pons | 340 | 170 | 170 |
| Pons | 15 | 8 | 7 |
| Pons | 105 | 55 | 50 |
| Pons | 47 | 23 | 24 |
| Pons | 521 | 251 | 270 |
| Pons | 2374 | 1191 | 1183 |
| Pons | 16 | 9 | 7 |
| Pons | 264 | 132 | 132 |
| Pons | 674 | 338 | 336 |
| Pons | 350 | 176 | 174 |
| Pons | 327 | 163 | 164 |
| Pons | 14 | 7 | 7 |
| Pons | 66 | 33 | 33 |
| Pons | 47 | 25 | 22 |
| Pons | 591 | 276 | 315 |
| Pons | 13 | 6 | 7 |
| Pons | 204 | 102 | 102 |
| Pons | 124 | 52 | 72 |
| Pons | 2334 | 1165 | 1169 |
| Pons | 83 | 36 | 47 |
| Pons | 29 | 15 | 14 |
| Pons | 49 | 24 | 25 |
| Medulla | 53 | 18 | 35 |
| Medulla | 610 | 302 | 308 |
| Medulla | 1031 | 514 | 517 |
| Medulla | 328 | 164 | 164 |
| Medulla | 81 | 40 | 41 |
| Medulla | 209 | 104 | 105 |
| Medulla | 153 | 77 | 76 |
| Medulla | 838 | 406 | 432 |
| Medulla | 1658 | 826 | 832 |
| Medulla | 1808 | 904 | 904 |
| Medulla | 1021 | 514 | 507 |
| Medulla | 101 | 51 | 50 |
| Medulla | 33 | 16 | 17 |
| Medulla | 931 | 467 | 464 |
| Medulla | 4 | 2 | 2 |
| Medulla | 27 | 14 | 13 |
| Medulla | 16 | 8 | 8 |

| | | | |
|---|---|---|---|
| Medulla | 168 | 87 | 81 |
| Medulla | 2606 | 1274 | 1332 |
| Medulla | 52 | 25 | 27 |
| Medulla | 486 | 243 | 243 |
| Medulla | 2766 | 1379 | 1387 |
| Medulla | 10 | 5 | 5 |
| Medulla | 65 | 34 | 31 |
| Medulla | 517 | 258 | 259 |
| Medulla | 59 | 29 | 30 |
| Medulla | 539 | 265 | 274 |
| Medulla | 1021 | 511 | 510 |
| Medulla | 897 | 450 | 447 |
| Medulla | 2331 | 1165 | 1166 |
| Medulla | 29 | 14 | 15 |
| Medulla | 247 | 125 | 122 |
| Medulla | 713 | 359 | 354 |
| Medulla | 35 | 18 | 17 |
| Medulla | 237 | 115 | 122 |
| Medulla | 99 | 48 | 51 |
| Medulla | 283 | 139 | 144 |
| Medulla | 1840 | 922 | 918 |
| Medulla | 761 | 379 | 382 |
| Medulla | 350 | 177 | 173 |
| Medulla | 56 | 27 | 29 |
| Medulla | 265 | 132 | 133 |
| Medulla | 28 | 14 | 14 |
| Medulla | 113 | 0 | 113 |
| Medulla | 67 | 0 | 67 |
| Medulla | 69 | 0 | 69 |
| Cerebellum | 126 | 55 | 71 |
| Cerebellum | 1336 | 630 | 706 |
| Cerebellum | 2742 | 1320 | 1422 |
| Cerebellum | 6777 | 3298 | 3479 |
| Cerebellum | 3334 | 1612 | 1722 |
| Cerebellum | 1030 | 498 | 532 |
| Cerebellum | 1233 | 590 | 643 |
| Cerebellum | 2162 | 1036 | 1126 |
| Cerebellum | 1503 | 721 | 782 |
| Cerebellum | 5709 | 2853 | 2856 |
| Cerebellum | 5660 | 2830 | 2830 |
| Cerebellum | 5118 | 2560 | 2558 |
| Cerebellum | 4866 | 2435 | 2431 |
| Cerebellum | 2490 | 1246 | 1244 |
| Cerebellum | 5742 | 2869 | 2873 |
| Cerebellum | 1310 | 658 | 652 |
| Cerebellum | 501 | 249 | 252 |
| Cerebellum | 348 | 147 | 201 |
| Cerebellum | 331 | 167 | 164 |
| Cerebellum | 87 | 46 | 41 |
