## Supplementary material for "Network structure of the mouse brain connectome with voxel resolution": Table S3

| ACRO | NAME | MACRO |
|---|---|---|
| MOp | Primary motor area | Isocortex |
| MOs | Secondary motor area | Isocortex |
| SSp-n | Primary somatosensory area, nose | Isocortex |
| SSp-bfd | Primary somatosensory area, barrel field | Isocortex |
| SSp-ll | Primary somatosensory area, lower limb | Isocortex |
| SSp-m | Primary somatosensory area, mouth | Isocortex |
| SSp-ul | Primary somatosensory area, upper limb | Isocortex |
| SSp-tr | Primary somatosensory area, trunk | Isocortex |
| SSp-un | Primary somatosensory area, unassigned | Isocortex |
| SSs | Supplemental somatosensory area | Isocortex |
| GU | Gustatory areas | Isocortex |
| VISC | Visceral area | Isocortex |
| AUDd | Dorsal auditory area | Isocortex |
| AUDp | Primary auditory area | Isocortex |
| AUDpo | Posterior auditory area | Isocortex |
| AUDv | Ventral auditory area | Isocortex |
| VISal | Anterolateral visual area | Isocortex |
| VISam | Anteromedial visual area | Isocortex |
| VISl | Lateral visual area | Isocortex |
| VISp | Primary visual area | Isocortex |
| VISpl | Posterolateral visual area | Isocortex |

| | | |
|---|---|---|
| VISpm | posteromedial visual area | Isocortex |
| VISli | Laterointermediate area | Isocortex |
| VISpor | Postrhinal area | Isocortex |
| ACAd | Anterior cingulate area, dorsal part | Isocortex |
| ACAv | Anterior cingulate area, ventral part | Isocortex |
| PL | Prelimbic area | Isocortex |
| ILA | Infralimbic area | Isocortex |
| ORB | Orbital area | Isocortex |
| AId | Agranular insular area, dorsal part | Isocortex |
| AIp | Agranular insular area, posterior part | Isocortex |
| AIv | Agranular insular area, ventral part | Isocortex |
| RSPagl | Retrosplenial area, lateral agranular part | Isocortex |
| RSPd | Retrosplenial area, dorsal part | Isocortex |
| RSPv | Retrosplenial area, ventral part | Isocortex |
| PTLp | or parietal associatio | Isocortex |
| TEa | nporal association ar | Isocortex |
| PERI | Perirhinal area | Isocortex |
| ECT | Ectorhinal area | Isocortex |
| MOB | Main olfactory bulb | Olfactory Areas |
| AOB | Accessory olfactory bulb | Olfactory Areas |
| AON | Anterior olfactory nucleus | Olfactory Areas |
| TT | Taenia tecta | Olfactory Areas |
| DP | Dorsal peduncular area | Olfactory Areas |
| PIR | Piriform area | Olfactory Areas |
| NLOT | Nucleus of the lateral olfactory tract | Olfactory Areas |
| COA | Cortical amygdalar area | Olfactory Areas |
| PAA | Piriform-amygdalar area | Olfactory Areas |
| CA | Ammon's horn | Hippocampal formation |

| | | |
|---|---|---|
| DG | Dentate gyrus | Hippocampal formation |
| ENT | Entorhinal area | Hippocampal formation |
| PAR | Parasubiculum | Hippocampal formation |
| POST | Postsubiculum | Hippocampal formation |
| PRE | Presubiculum | Hippocampal formation |
| SUB | Subiculum | Hippocampal formation |
| ProS | Prosubiculum | Hippocampal formation |
| HATA | Hippocampo-amygdalar transition area | Hippocampal formation |
| APr | Area prostriata | Hippocampal formation |
| CLA | Claustrum | Cortical subplate |
| EP | Endopiriform nucleus | Cortical subplate |
| LA | Lateral amygdalar nucleus | Cortical subplate |
| BLA | Basolateral amygdalar nucleus | Cortical subplate |
| BMA | Basomedial amygdalar nucleus | Cortical subplate |
| PA | Posterior amygdalar nucleus | Cortical subplate |
| STRd | Striatum dorsal region | Striatum |
| STRv | Striatum ventral region | Striatum |
| LSX | Lateral septal complex | Striatum |
| sAMY | Striatum-like amygdalar nuclei | Striatum |
| PALd | Pallidum, dorsal region | Pallidum |
| PALv | Pallidum, ventral region | Pallidum |
| PALm | Pallidum, medial region | Pallidum |
| PALc | Pallidum, caudal region | Pallidum |
| DORsm | Thalamus, sensory-motor cortex related | Thalamus |

| | | |
|---|---|---|
| DORpm | Thalamus, polymodal association cortex related | Thalamus |
| PVZ | Periventricular zone | Hypothalamus |
| PVR | Periventricular region | Hypothalamus |
| MEZ | Hypothalamic medial zone | Hypothalamus |
| LZ | Hypothalamic lateral zone | Hypothalamus |
| MBsen | Midbrain, sensory related | Midbrain |
| MBmot | Midbrain, motor related | Midbrain |
| MBsta | Midbrain, behavioral state related | Midbrain |
| P-sen | Pons, sensory related | Pons |
| P-mot | Pons, motor related | Pons |
| P-sat | Pons, behavioral state related | Pons |
| MY-sen | Medulla, sensory related | Medulla |
| MY-mot | Medulla, motor related | Medulla |
| CBX | Cerebellar cortex | Cerebellum |
| CBN | Cerebellar nuclei | Cerebellum |
